# A hippocampal population code for rapid generalization

**DOI:** 10.1101/2025.03.15.643456

**Authors:** Wenbo Tang, Hongyu Chang, Can Liu, Salma Perez-Hernandez, William Y. Zheng, Jaehyo Park, Azahara Oliva, Antonio Fernandez-Ruiz

## Abstract

Generalizing from experience and applying prior knowledge to new situations is essential for intelligent behavior. Traditional models attribute this generalization to gradual statistical learning in the neocortex. However, such a slow process cannot account for animals’ rapid generalization from limited experience. Here, we demonstrate that the hippocampus supports rapid generalization in mice by generating disentangled memory representations, where different aspects of experience are encoded independently. This code enabled the transfer of prior knowledge to solve new tasks. We identify specific circuit mechanisms underlying this rapid generalization. We show that the seemingly random changes in individual neuronal activity over time and across environments result from structured circuit-level processes, governed by the dynamics of local inhibition and cross-regional cell assemblies, respectively. Our findings provide computational and mechanistic insights into how the geometric structure and underlying circuit organization of hippocampal population dynamics facilitate both memory discrimination and generalization, enabling efficient and flexible learning.

## Introduction

Learning requires that animals balance two opposing cognitive demands: discriminating between related but distinct experiences and generalizing common elements across those experiences. While discrimination is important for encoding new information without interference with previously stored knowledge ^1-6^, generalization enables making inferences in novel situations, integrating individual experiences into overarching knowledge structures, and learning abstract concepts ^7-13^. It is, therefore, a key feature of intelligent behavior. Yet, how the brain achieves this after only limited experience is not understood.

Because memory generalization comes at the expense of discrimination, a proposed solution to this dilemma has been to assign each of these functions to a different brain structure. In this framework, the hippocampus supports initial learning and discrimination of episodic memories, while the neocortex gradually develops generalized representations integrating over multiple experiences ^9,12,14-16^. While this segregation of functions has received extensive support ^10,12,17-20^, it cannot account for recent observations on the role of the hippocampus in rapid inference, concept learning and generalization ^7,8,21-24^. An alternative possibility is that the initial hippocampal encoding of episodic memories involves yet-to-be discovered mechanisms that facilitate both rapid generalization and discrimination.

Previous work has shown that the hippocampus combines information about the location, time and content of each experience into unified “conjunctive” representations ^10,25-30^. Hippocampal representations of different experiences, even if sharing common features, tend to be largely uncorrelated (Figures 1A and 1B) ^4,5,31-38^. Conjunctive and uncorrelated representations support memory specificity and discrimination, but they do so at the cost of generalizability. For instance, if eating in different dining halls is encoded as unique and uncorrelated representations, how can the brain immediately recognize and exploit a shared structure upon entering a new one (e.g., starting at the beginning of the tray line before heading to the cashier)? Recent theoretical work on how artificial neural networks achieve generalization offers new insights into this question ^39-41^. Encoding different aspects of experience as “disentangled” representations – where different features are split into independent dimensions of neural population activity, rather than being encoded as a unified representation – allows to re-combine them later to represent novel situations (Figure 1B, *right*). In the previous example, if, at the time of visiting different dining halls, the food eaten and the hall’s spatial layout are encoded in a disentangled manner, the brain can readily re-use the layout representation to recognize a similar structure in a new dining hall. Such rapid generalization of memory representations can support flexible behavior, such as efficiently navigating a new dining hall. Here, we found that hippocampal population dynamics enable both efficient memory discrimination as well as generalization, supporting rapid learning based on the generalization of prior knowledge.

**Figure 1.**
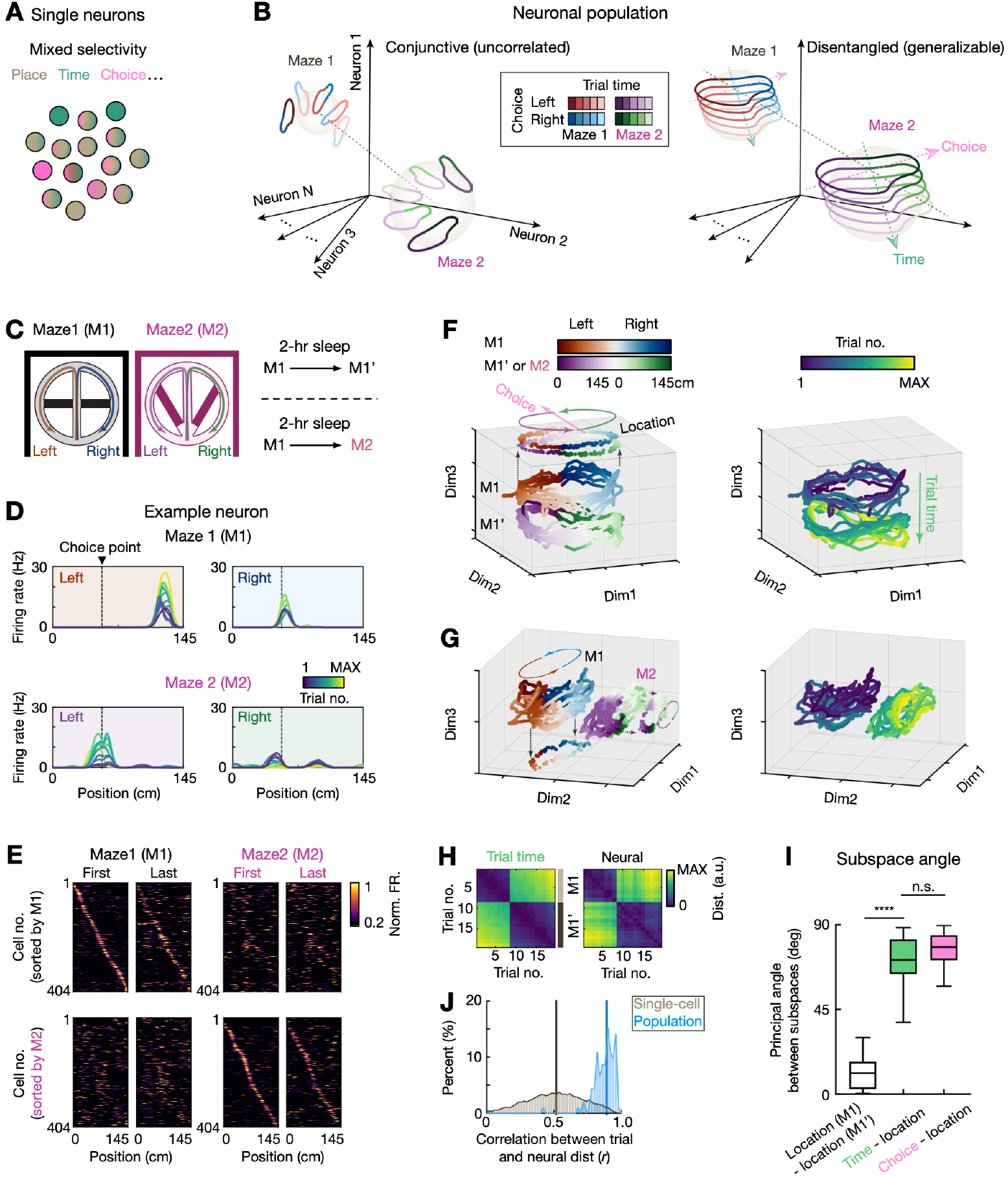
Disentangled coding of task variables by hippocampal population dynamics. (**A**) Single hippocampal neurons show mixed selectivity for places, time and choices. (**B**) The geometry of hippocampal representations (i.e., the arrangement of *N*-dimensional neural activity vectors) can be either conjunctive (*left*) or disentangled (*right*). *Left:* Conjunctive representations are characterized by distinct neural trajectories for each single experience, which are uncorrelated among different experiences. *Right:* In disentangled representations the coding direction (arrow headed lines) for each variable is independent (i.e., orthogonal) of coding of the others, and can be generalized across different experiences. (**C**), Mice performed delayed alternation (left - L - vs. right - R -) in two different mazes (M1 and M2; M1’, re-exposure to M1). (**D**) Example CA1 neuron showed conjunctive selectivity for locations, trial time and choices (L vs R), but this selectivity did not generalize across mazes. Ordinal response to trial number was apparent for the L-trajectory in M1, but not in M2; choice predictive firing preceding the decision point (vertical dashed line) was stronger for R than L in M1 but not in M2. **e**, Normalized rate maps of all CA1 spatial-tuned neurons during the first and last trial in M1 and M2, sorted by peak activity in M1 (*top*) and M2 (*bottom*). **(F** and **G)** 3D embeddings of neural manifolds using unsupervised UMAP for two sessions in M1 **(F)**, or in M1 and M2, respectively **(G)**. Each dot represents population activity in a 200-ms bin, color coded by locations and choice types (*left*) or trial numbers (*right*). Trial-averaged activity was projected on a 2-D plane for ease of visualization of ring structure (see also Figure S2A). **h**, Representational dissimilarity matrix (RDM) quantified as the Euclidean distance between the median time stamps (*left*) and neuronal population vectors (*right*) of each pair of trials in the same maze (M1 and M1’ session). (**I**) Correlation between the RDMs of trial time and neural activity using single cells (gray) or population (blue) (\*\*\*\**p* = 1.32e-56, Kolmogorov-Smirnov test comparing the single-cell versus population distribution). Vertical lines on histograms: median. (**J**) Principal angles between different coding subspaces. The 2-dimensional subspace in the *N*-dimensional neural state space that contains the most variance of one variable was identified as the coding subspace of that variable (Figure S2B). *From left to right*, principal angles between M1 and M1’ location subspaces, between time vs. location subspaces and between choice vs. location subspaces on the same maze (*n* = 20 session pairs; \*\*\*\**p* = 4.05e-10, n.s., *p* = 0.42, Kruskal-Wallis test with Dunn’s *post hoc*). Note that quantifications in **(H-J)** were performed in the *N*-dimensional neural state space, rather than the reduced UMAP space. See also **Figures S1** and **S2**.

### The disentangled structure of the hippocampal memory code

We implanted mice with ultra-high-density active silicon probes allowing us to simultaneously record up to ∼400 CA1 cells while animals performed the same hippocampus-dependent memory task (delayed spatial alternation) in different mazes. In this task, mice need to correctly choose the previously unvisited arm after a delay between two upcoming options (the left or right arm). The different mazes shared a similar structure (resembling a figure 8) but largely differed in both local and global cues (Figure 1C). This behavioral paradigm enabled us to investigate both memory discrimination (of the different mazes) and generalization (of the common task structure). We recorded mice (*n* = 6) in morning and afternoon sessions either in the same or different mazes, interleaved with 2-hour rest in the home cage (Figure 1C). We first analyzed how individual cells encoded the three key variables in this task: location, time (trial order) and choice (upcoming left vs. right turn). We observed a heterogeneity of single neuron responses, with some neurons preferentially encoding one of the variables and many others tuned to all three (Figures 1D and S1). This “mixed selectivity” of hippocampal cells (Figures 1A and S1) is indicative of a conjunctive code ^42,43^. Notably, single neuron tuning properties did not generalize across mazes (Figures S1E-S1H). That is, a neuron encoding the left choice in one maze could be weakly tuned to choices, but strongly selective to locations or time in the other maze (Figures 1D and S1A). At the ensemble level, the representation of the two mazes was completely uncorrelated, indicative of “global remapping” ^3,34,44^ (Figures 1E and S1F-S1H). In contrast, ensemble responses for morning and afternoon sessions in the same maze (and between early and late trials within a session) were highly correlated, albeit with varying degrees of temporal “drift” in neuronal tuning (i.e., changing response to the same location over trials) ^38,45,46^ across the two sessions (Figures 1D, 1E and S1). Therefore, changes in neuronal tuning to both time and space were seemingly random and uncorrelated (Figure S1), in agreement with previous reports ^33,34,45,46^. These properties of the hippocampal code offer a good substrate for discriminating related experiences (similar mazes) but pose a challenge for identifying common structural elements across them.

To test whether hippocampal representations for different experiences were encoded in a conjunctive or disentangled manner (Figure 1B), we first visualized the geometric structure of hippocampal population activity by embedding it in a 3D space using unsupervised Uniform Manifold Approximation and Projection (UMAP). The structure of population activity in UMAP space resembled the topological structure of the maze (a ring connecting left and right trajectories in succession; Figures 1F, 1G and S2A) ^11,47-51^.

Furthermore, although UMAP was calculated in a fully unsupervised manner without including temporal information, we observed that successive laps appeared as adjacent parallel trajectories in the manifold, with a larger gap between two sessions separated by a 2-hour rest (Figure1F). Importantly, location, time and choice were encoded along three different dimensions, largely orthogonal, within the manifold (Figures 1F and S2A). The same procedure applied to days when mice ran in two different mazes revealed that neural trajectories in both mazes lay in different subspaces that appeared rotated with respect to each other (Figures 1G and S2A). These results suggest a disentangled code in the mouse hippocampus, a property previously thought to primarily characterize cortical representations ^11,52-55^. Due to the limitations of UMAP in preserving global data structure, we analyzed the geometry of hippocampal representations in the original high-dimensional neural state space. First, we confirmed that the trial-by-trial drift in population activity was strongly correlated with time, instantiating a time code (Figure 1H) ^46,51,56^. This property largely emerged at the population level, while single neurons showed varying changes of place-field tuning drift over time (Figures 1I and S1). To confirm that location, time and choice were encoded in a disentangled manner, we calculated the principal angle between coding subspaces of different variables ^55,57,58^. Two variables are disentangled if their coding subspaces are approximately orthogonal, as they can be independently readout by a linear decoder ^7,43,54^. We found angles close to 90 degrees between location, time and choice coding subspaces (Figure 1J) with minimal overlap between one another (Figures S2B-S2G), supporting a disentangled coding scheme.

### Disentangled hippocampal dynamics support memory generalization

We hypothesized that the disentangled population code we found facilitates generalizing common features across different experiences to support efficient learning. To test this, we examined hippocampal representations as mice learned to perform the same delayed alternation task in a sequential manner across three different mazes (Figure 2A). Mice were trained in the familiar maze (M1) until they reached an average 80% (5-6 days) of correct choices per session (83.9 ± 3.2%, *n* = 5 mice). Behavioral performance dropped significantly the first time animals were placed in the second maze (M2), but increased rapidly the following day (Figure 2B). Furthermore, the first time mice experienced M3, performance was significantly higher, at the same level as the familiar M1 (Figure 2B). This result suggests that mice were using the knowledge of the task acquired in one maze to effectively solve the same problem in a different one (“transfer learning”) ^8,10,59^.

**Figure 2.**
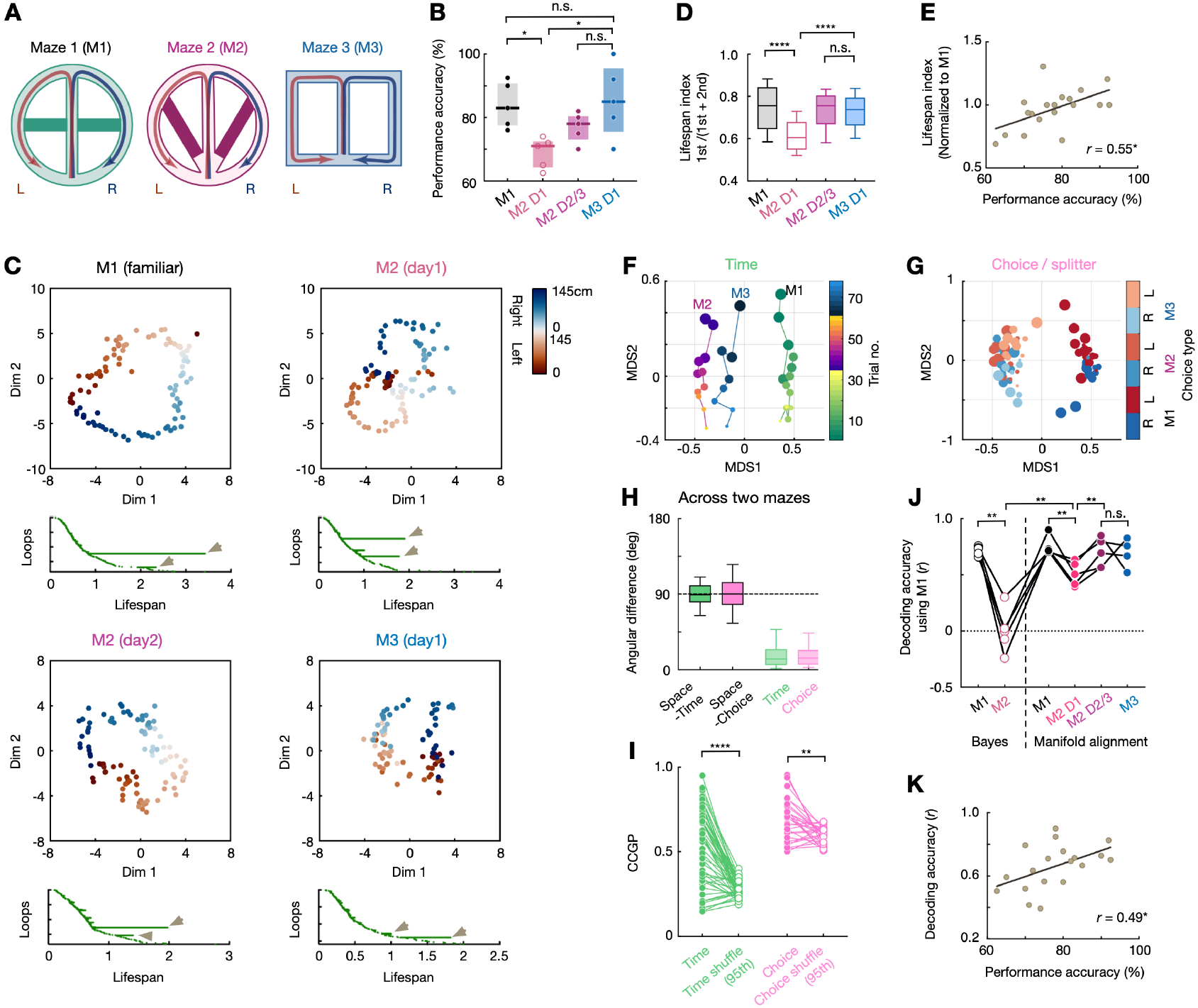
Hippocampal population dynamics support representational and behavioral generalization. (**A**) Schematic of experimental setup. (**B**) Behavioral performance accuracy for the last M1 session, the first and later M2 sessions, and the first M3 session, respectively (from left to right, \**p* = 0.012, 0.048, n.s., *p* = 0.98 and 0.71, one-way ANOVA test with Tukey’s post hoc). D1: day 1; D2/3: day 2 or 3. (**C**) *Top*, visualization of the topological structure of neural manifolds during the last session on M1, the first and second session on M2 and the first session on M3. *Bottom*, related persistent co-homology lifespan of 1-dimensional feature (*H1*, or loops). Arrows show 2 most prominent bars (or loops). Note the transformation of the neural manifold from 2 loops to 1 loop with learning. (**D**) Lifespan index, measured as the bar length ratio between the first and second most prominent loops (\*\*\*\**p* < 1e-16, n.s., *p* = 0.44, Kruskal-Wallis test with Dunn’s *post hoc*; same sessions used in **B** and **C**). (**E**) Correlation between behavioral performance accuracy and lifespan index (normalized to M1 for each animal; \**p* = 0.032, *r* = 0.55, Pearson correlation). **(F** and **G)** Structure of population activity similarity across trial numbers **(F)** and choice types **(G)** on the three mazes embedded in a 2D space via MDS. Dot size with color code indicates trial number. Note that coding direction of trial time (solid lines) and the coding direction of spatial contexts (M1-M2-M3) were nearly orthogonal in the embedding shown in **(F)**. Similar orthogonality of choices and spatial contexts is shown in **(G)**. (**H**) From left to right, angular difference between the coding direction of spatial contexts and temporal drift, the direction of spatial contexts and choice, the direction of temporal drift and choice between two spatial contexts (i.e., two mazes). While space and time dimensions were orthogonal, the coding direction of time and choice were consistent across different mazes (*n* = 42 session pairs from 5 mice). (**I**) CCGP for time and choice decoding across mazes (*n* = 2 choice x 27 session pairs for time, and 27 session pairs for choice from 5 mice, each paired line for one pair; \**p* = 0.014, \*\**p* = 0.003, Wilcoxon paired test, compared to the 95th percentile of shuffled values). (**J**) Location decoding accuracy using the decoder built on M1 for prediction of M2 D1, M2 D2/3, and M3. First two columns are from canonical Bayesian decoders, and the last four columns are from manifold alignment (from left to right, \*\**p* = 0.0093, 0.0012, 0.0098, 0.0019, *n* = 5 mice; n.s., *p* = 0.77, *n* = 4 mice; Paired *t*-tests). (**K**), Correlation between behavioral performance accuracy and cross-maze decoding accuracy with manifold alignment (\**p* = 0.014, *r* = 0.49, Pearson correlation). See also **Figure S3**.

To examine the learning-dependent evolution of hippocampal manifold geometry, we compared its topological structure across days and mazes using persistent co-homology ^49,60,61^. The neural manifold had a ring shape (single-loop topology) in the familiar M1 (Figure 2C, *upper left*), suggesting that mice had learned the latent structure of the task (i.e., running through the central stem in right versus left-bound journeys correspond to two different task states). During the initial session in M2, the hippocampal manifold resembled instead an 8 (two-loop topology) (Figure 2C, *upper right*). This difference in shape suggests that mice did not immediately recognize the shared task structure of the two mazes, as supported by the drop in their performance (Figure 2B). Indeed, the hippocampal manifold became a ring on the second day in M2 and readily displayed a ring topology on the first day in M3 (Figure 2C, *bottom*), mirroring the generalization of behavioral performance across mazes (Figure 2D). Furthermore, we found a significant positive correlation between manifold topology and behavioral performance at the individual session level (Figure 2E). These changes in manifold topology with learning were paralleled by corresponding changes in the proportion of choice-selective cells on the center stem (i.e., splitter cells; Figure S3A). Notably, such topological change cannot be simply explained by familiarization, animals’ locomotion, or physical layouts of the mazes, but reflected the learning of a latent task’s structure, as the neural manifolds during a forced alternation task (thus without memory demands) on the same maze consistently expressed a two-loop structure (Figures S3B-S3C).

An additional requisite for disentangled neural representations to support task generalization is that the encoding of a given variable must be consistent across different mazes. To test this prediction, we compared the similarity in the representational structure of hippocampal population activity across mazes by embedding it into a common 2D space using multidimensional scaling (MDS) ^62^. We found a shared coding dimension for time (trial progression) across all three mazes (Figures 2F and 2H), as well as a common coding dimension for choice (Figures 2G and 2H). Consistent with our subspace analysis (Figure 1J), the coding dimensions for time and choice were orthogonal to the coding dimension for space (Figure 2H). We further confirmed this generalizability using a decoding approach. We trained a linear classifier to discriminate trial numbers or choice types in one maze and tested its performance in the other two mazes (cross-conditional generalization performance -CCGP-; Figures S3D-S3E) ^7,8^. We observed a significant generalized decoding performance across mazes for the two variables (Figure 2I). These results indicate that the disentangled population coding of distinct task variables in the hippocampus generalizes across different environments, confirming a key prediction of our hypothesis (Figure 1B).

We further reasoned that a generalizable hippocampal code could capture the shared latent structure of the task across mazes despite the global remapping of individual place cells (Figure 1E). While the neural manifolds for two mazes appeared rotated with respect to each other, we were able to align them by applying a rigid geometric transformation (Figure S3F) ^63^. Next, we trained a linear classifier to decode spatial locations using the manifold of one maze and tested it on data from the other mazes (from same-day recordings), before and after manifold alignment. Cross-maze spatial decoding accuracy was below chance level before the alignment but significantly increased after applying the transformation (Figures 2J and S3G). Spatial decoding performance after cross-maze manifold alignment displayed a similar evolution over days as learning performance (Figure 2J), that is, it was lower in the first session in M2 but increased in the second one and it was already high in the first session in M3. Indeed, the cross-maze decoding accuracy was significantly correlated with behavioral performance of each session (Figure 2K). Therefore, the decoding performance after manifold alignment did not only reflect the shared maze geometry but it was also influenced by behavioral performance. Altogether, these results suggest that a generalizable latent manifold structure shared across different mazes emerged in the hippocampus after task learning.

### Cell assembly correlations constrain remapping dynamics across environments

We found that the hippocampus encodes different experiences using a disentangled population code (Figure 1), and that the representations of individual task variables generalize across different environments (Figure 2). We next investigated the physiological mechanisms that support these two key features of hippocampal representations. We made two predictions. 1) The critical feature that enabled generalization is that coding dimensions of task variables remained consistent across mazes. How is this possible when single-unit responses appear to change randomly (remap) across environments? We postulate that, in contrast to the common assumption in the field ^2,3,33,34^ (but see ^28,64^), place cell remapping in a new environment is not completely random but constrained by a dedicated circuit mechanism such that correlations across environments are preserved. 2) We hypothesize that disentangled representations of different task variables, such as time and location, are enabled by distinct circuit mechanisms supporting each of them. While individual cells show mixed selectivity for different task variables, such mechanistic dissociating would enable their independent representation at the population level.

Previous work has shown how precisely timed excitatory inputs drive the reconfiguration of CA1 place fields ^65-67^. We hypothesize that the underlying circuit organization and input synchrony that shapes the structure of hippocampal functional cell assemblies (groups of coactive cells with common synaptic inputs) ^68,69^ may preserve spatial correlations across environments. We first explored this hypothesis by conducting simulations in a simplified firing-rate model of the hippocampal network that included the main input region to CA1 place cells, CA3 (Figure 3A). The objective of this model framework is to optimize the output neural representation, so that the geometric mismatch between the input and the representation is minimized ^70^. The place cells in each layer receive excitatory feedforward inputs and compete via local inhibition. Synaptic noise in excitatory and inhibitory inputs cause representational drift and drive the network to explore the near-optimal space. We gave two different circular mazes as spatial inputs to the model (Figure 3A), and the model generated localized place fields in CA3 and CA1, which exhibited global remapping across mazes (Figure 3B). We then detected coordinated CA3-CA1 cell assemblies across the two regions by applying singular value decomposition (SVD) on the correlation matrix of cross-regional population activity only from the first maze (Figure S4A). Remarkably, the remapping of CA1 cells that were members of the same CA3-CA1 assembly was highly correlated across mazes, despite place-field remapping of the whole CA1 population being completely uncorrelated (Figures 3C-3E). This result suggests that the functional structure of CA3 inputs preserves correlations in CA1 remapping across environments.

**Figure 3.**
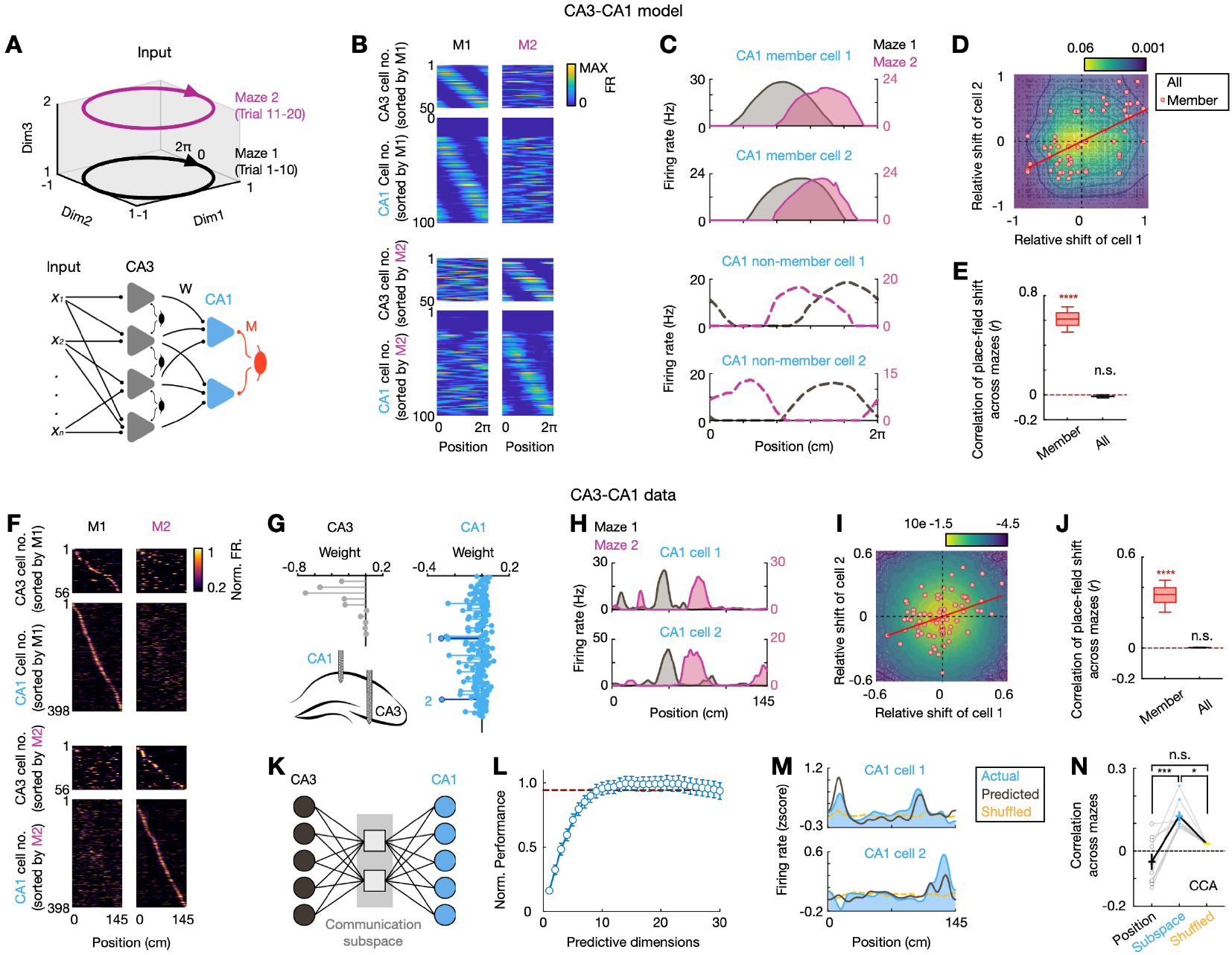
Coherent remapping across environments in CA3-CA1 assemblies. (**A**) CA3-CA1 model architecture. Spatial inputs from two circular mazes (*top*) were feedforward onto CA3. Both CA1 and CA3 pyramidal cells (triangles) received local lateral inhibition (ellipses). (**B**) Simulated CA3 and CA1 place-field activity, sorted by peak activity in M1 (*top*) and M2 (*bottom*). (**C**) Rate maps of the two CA1 member (*top*) and non-member (*bottom*) cells across Maze 1 (gray) and 2 (purple). Note the coherent remapping for member cells. (**D**) Place-field center-of-mass (COM) shift across Maze 1 and Maze 2 for CA1 cell pairs belonging to the same CA3-CA1 assemblies (red circles) versus all CA1 pairs (black dots; *n* = 4 model simulations; density map shows joint distribution of place-field shift for all CA1 pairs; red line, best linear fit for member pairs). (**E**) Correlation of place-field COM shifts across mazes for assembly members (red; *p* < 1e-16, signed rank test, *n* = 1000 permutations) and all pairs (black; *p* > 0.99). (**F**) Rate maps of simultaneously recorded CA3 and CA1 cells from experimental data. Data is shown as in **(B)**. **(G** and **H)** Example CA3-CA1 assembly from experimental data. **(G)** Assembly weights, with two CA1 member cells highlighted in dark blue. **(H)** Rate maps of the two member cells shown in **(G)**. (**I**) Place-field COM shift across Maze 1 and Maze 2 for all recorded CA1 pairs (*n* = 16 sessions from 3 mice). Data are presented as in **(D)**. (**J**) Correlation of place-field COM shifts across mazes (\*\*\*\**p* < 1e-16 for members, and *p* = 0. 044 for all pairs, signed rank test). (**K**) Schematic of CA3-CA1 communication subspace. (**L**) Number of predictive dimensions needed to achieve full predictive performance was 7 ± 3 (mean ± SD; 10 sessions from 4 mice). Error bars indicate standard errors (SEs) via cross-validation. Red dashed line as (mean - SE) of peak performance. (**M**) Example CA1 cells’ spatial firing rate maps in M2 (blue), compared with the firing rate predicted from CA3-CA1 communication subspace in M1 (black) and from temporally shuffled M2 CA3 activity (yellow). (**N**) Correlation of CA3-CA1 activity across two mazes aligned to their communication subspaces was significantly higher than that from their temporally shuffled data (\**p* = 0.011), and from position alignment (\*\*\**p* = 0.0004, *n* = 10 session pairs from 3 mice, Friedman test with Dunn’s *post hoc*). See also **Figure S4**.

Guided by the model prediction, we conducted new neural recordings simultaneously in CA1 and CA3 areas while mice performed the same alternation task across two different figure-8 mazes (Figures 3F-3G and S4B). To test whether the structure of cross-regional cell assemblies preserves remapping correlations across environments, we used the same method as with the simulated data. We detected cell assemblies of coactive CA3 and CA1 cells in one maze and examined spatial correlations between mazes (Figures 3G-3J and S4C-S4G). Consistent with our model prediction, while random remapping was observed for all CA1 cell pairs (Figures 3I-3J and S4D-S4E), as previously reported ^31,33,34,37,38,44^, we found that remapping of CA1 cell members of the same CA3-CA1 assemblies was highly correlated between mazes (Figures 3G-3J and S4D-S4G). That is, for two assembly members, the place-field shift of one member between M1 and M2 was similar to the shift observed for another member (note that assemblies were defined by coactivity patterns in M1 exclusively). Interestingly, when assemblies were detected as groups of co-active CA1 cells, without considering CA3, correlation of assembly members across mazes were significantly lower (Figures S4F-S4G), underscoring the key role of shared CA3 inputs in preserving spatial correlations. Overall, our simulation and experimental results demonstrate that hippocampal remapping is not a completely random process but heavily constrained by the structure of assembly dynamics that is brought about by the functional organization of the CA3-CA1 network.

To further verify that CA3-CA1 coordination preserves functional correlations across environments, we employed a complementary analytical approach. We used dimensionality reduction methods to identify a specific subspace in the CA3-CA1 high-dimensional state space where effective coordination between these two regions took place (i.e., “communication subspace”; Figure 3K) ^71,72^. Applying this method to CA3-CA1 spike trains from M1, we found a low-dimensional communication subspace between these two regions (Figure 3L). Remarkably, the spatially modulated activity in the subspace from one maze was significantly predictive of the spatially modulated activity in a different maze (Figure 3M), suggesting a consistent alignment of the communication subspaces across environments. Indeed, the communication subspaces independently identified for M1 and M2 were consistently aligned, significantly better than CA3-CA1 temporally shuffled data (Figures 3N and S4H). In contrast, using the communication subspaces defined by the CA1 spatial responses across the two mazes, instead of CA3-CA1 coordination, resulted in a poor alignment (Figure 3N), consistent with the observed uncorrelated spatial remapping of the whole CA1 population (Figures 3I-3J). This result indicates that the low-dimensional population structure of CA3-CA1 interactions was preserved across mazes despite the global remapping of their single cell place fields.

### Distinct circuit mechanisms support spatial remapping and temporal drift

We next investigated whether the disentanglement of representations for spatial location and temporal progression by CA1 population activity is mediated by distinct circuit mechanisms supporting each of them. We first explored this issue with the CA1 firing-rate model (Figure 4A). The model generated localized CA1 place fields drifting across trials in the same circular maze (Figure 4B). Given our results (Figure 3) and previous work ^65-67^ on the critical role of CA3 inputs in CA1 place cell expression and remapping, we examined the effect of their disruption in the model. Partial silencing on CA3 inputs in the model (Figure 4A) resulted in CA1 place field remapping within the same maze (Figure 4A-4C). At the population level, this remapping manifested as a rotation of the neural manifold of ∼90 degrees (Figures 4A and S5A). These results were similar to our experimental observations of hippocampal remapping across different mazes (Figure 1). In contrast, the temporal structure of trial-by-trial drift was preserved (Figures 4B and 4D), suggesting that distinct circuit mechanisms may underlie the two phenomena.

**Figure 4.**
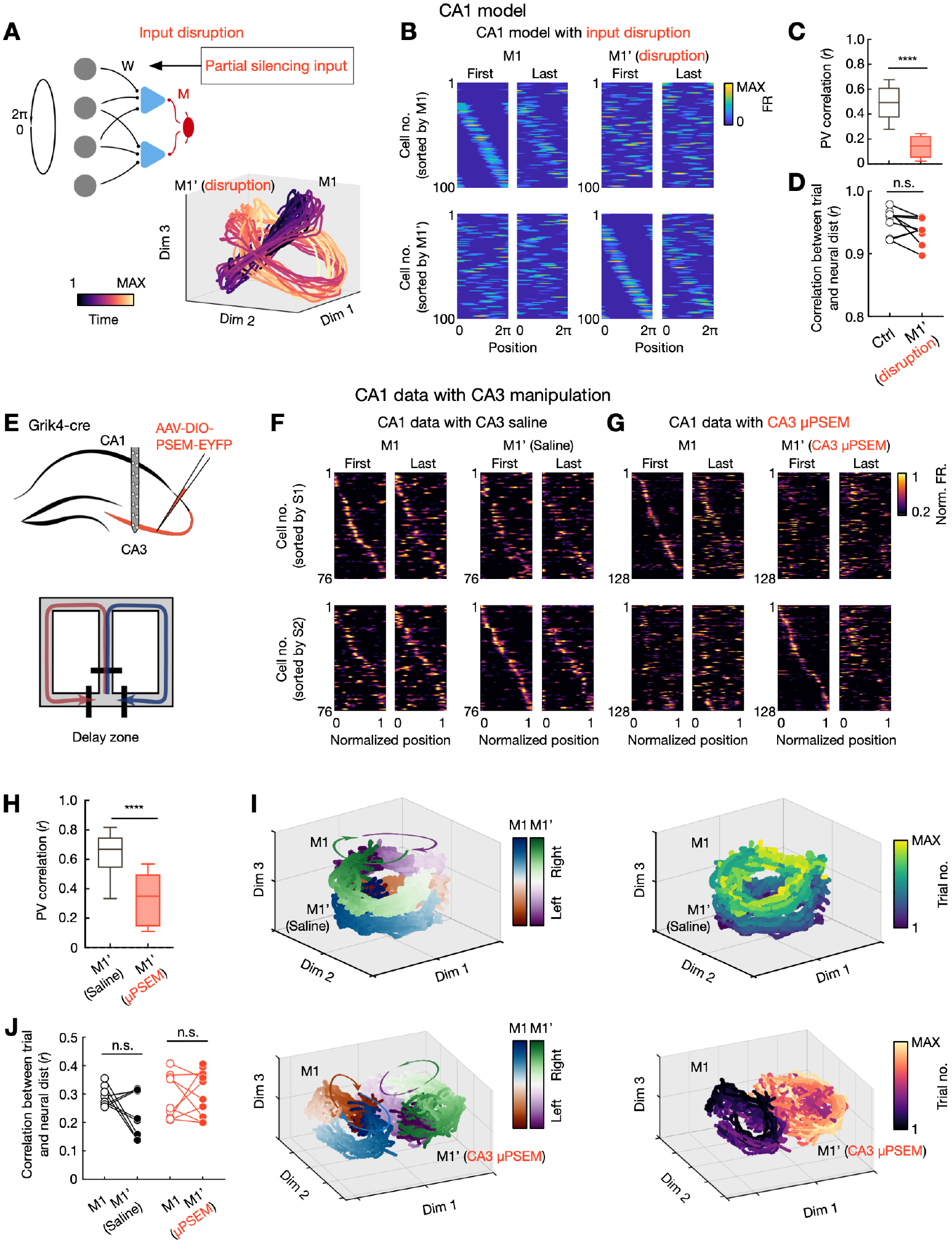
Perturbation of CA3 inputs induced global remapping in CA1, but did not affect temporal drift. (**A**) Disrupting feedforward inputs (reducing their strength and enhancing synaptic noise) in the CA1 model (*left*), induced a manifold rotation in the same maze (*right*; early vs late trials). (**B**) Simulated place-field activity before (M1) and during (M1’) input disruption on the same circular maze, sorted by the peak firing location during M1 (*top*) and M1’ (*bottom*) trial blocks. Note global remapping. (**C**) Population vector (PV) correlation across M1 and M1’ from the model simulations (*left*; *n* = 9 simulations, \*\*\*\**p* < 1e-16, rank-sum test). (**D**) Input disruption did not affect the representational drift (correlation between trial time and neural distances; n.s., *p* = 0.13, Wilcoxon paired test). (**E**) Schematic of CA1 recording with partial CA3 silencing (*top*). Mice (*n* = 4) performed a delayed alternation task on a figure-8 maze (*bottom*). **(F** and **G)** Rate maps of CA1 cells from an example control **(F)** and CA3 disruption **(G)** session. Data are shown as in **(B)**. Note the remapping during re-exposure to the same maze. (**H**) Population vector (PV) correlation across M1 and M1’ from the experimental data (*right*; \*\*\**p* =0.0008, *n* = 10 and 11 sessions for saline and μPSEM conditions respectively, rank-sum test), similar to that from the model **(C)**. (**I**) Manifold dynamics during control (*top*) and CA3 disruption (*bottom*) sessions. Note the parallel representational drift with time in the control condition, similar to Figure 1F, and the rotational dynamic with CA3 perturbation, similar to the remapping shown Figure 1G. (**J**) CA3 perturbation did not affect CA1 representational drift with time. Data are presented as in **(D)**. Each paired line for one session pair (black, control sessions, n.s., *p* = 0.25, *n* = 10 session pairs red, experimental sessions; n.s., *p* = 0.63, *n* = 9 session pairs, Wilcoxon paired test, n = 4 mice). See also **Figure S5**.

To experimentally validate the model’s prediction, we analyzed a previous dataset ^73^, in which CA3 was perturbed using a chemogenetic silencing approach (PSAM-GlyR) ^74^, while simultaneously recording CA1 (Figure 4E). We compared CA1 population dynamics in consecutive control and manipulation sessions (systemic injection of saline versus the PSAM-GlyR agonist μPSEM792), while mice performed a figure-8 delayed alternation task (Figure 4E). This perturbation partially silenced CA3 with reduced firing rate and reduced activity correlation among its pyramidal cells (Figure S5B), mirroring the model manipulation (Figure 4A), while it preserved basic CA1 place field features, albeit with a mild reduction in firing rates ^73^. Importantly, CA3 perturbation (but not a control vehicle injection) induced global remapping in CA1, manifested both as the decorrelation of spatial maps for individual cells (Figures 4F-4H) and as a manifold rotation (Figures 4I and S5B), consistent with our simulation results. In contrast, the structure of representational drift and its correlation with time were preserved (Figures 4I and 4J), supporting a mechanistic dissociation between place cell temporal drift and spatial remapping. Optogenetic perturbations of another major input to CA1, the medial entorhinal cortex (MEC), similarly induced global remapping while preserving temporal drift (Figures S5C-S5G). These results suggest that CA3 and MEC inputs determine CA1 spatial remapping dynamics but do not play a major role in regulating the structure of temporal drift.

We next sought to identify the mechanism regulating the ordered structure of place cell temporal drift. In addition to feedforward excitatory inputs, another important circuit element contributing to CA1 dynamics are inhibitory inputs from local interneurons ^75-78^. We reasoned that, beyond a broad control of pyramidal cell excitability, temporarily structured inhibition may play an additional role in regulating place cells’ spatio-temporal dynamics.

To perturb the temporal structure of inhibition in the model, we injected white noise into all CA1 inhibitory synapses (Figure 5A). This manipulation increased the temporal variability of interneuron activation and enhanced their synchronization at each moment. Place field properties were preserved by the manipulation, but their stability (trial-by-trial correlation) increased (Figures 5B-5D), suggesting a reduction in place cell temporal drift. This alteration was also manifested as a disruption in the ordered progression of trial-by-trial trajectories in the neural manifold (Figure S6A). The manipulation did not induce place field remapping (Figures 5B-5D), suggesting a specific contribution of the temporal structure of local inhibition to hippocampal representational drift.

**Figure 5.**
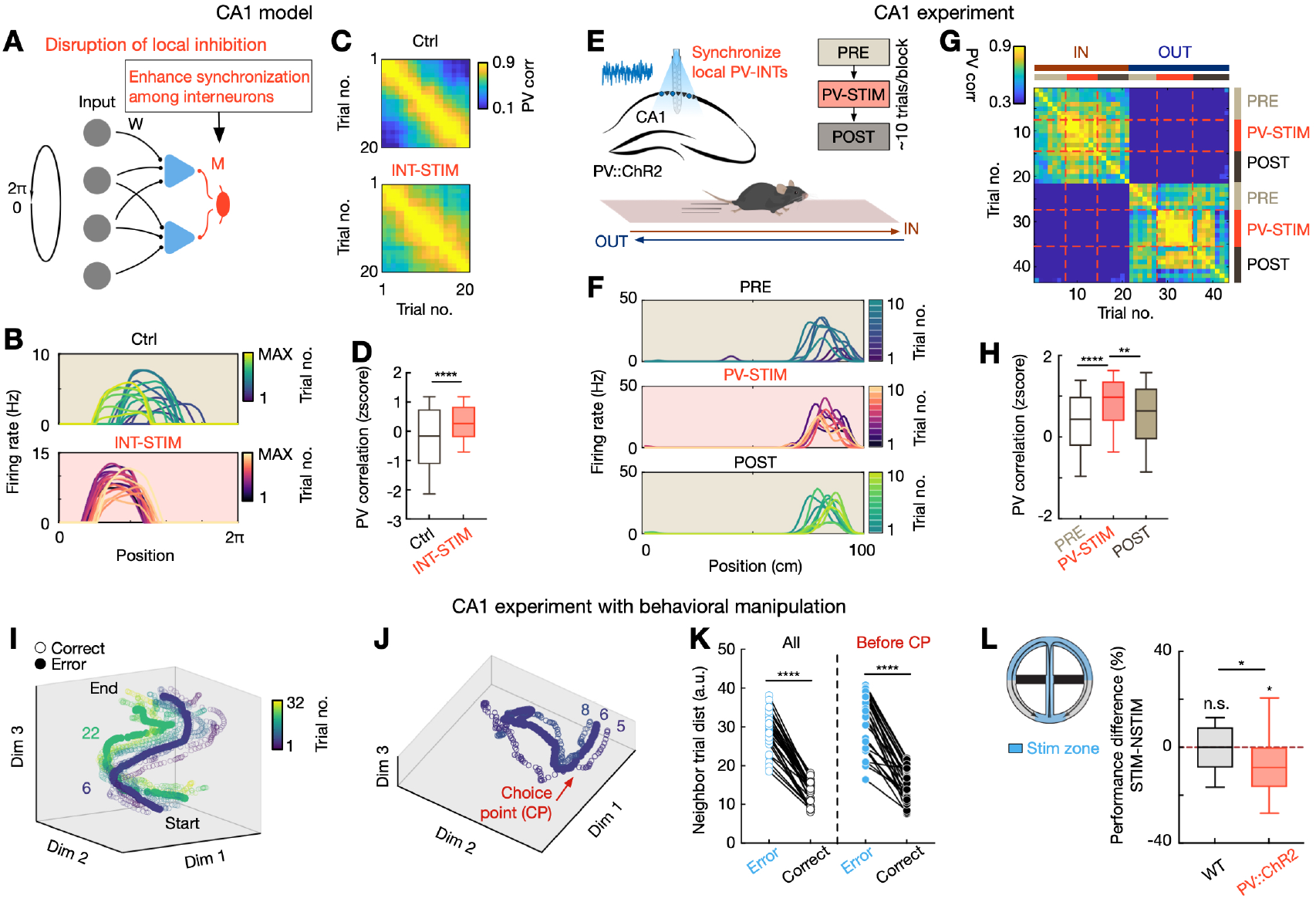
Lateral inhibition by CA1 PV interneurons promotes representational drift. (**A**) Model schematic. (**B**) Example simulated place cell in control condition (*top*) and with the manipulation of lateral inhibition (INT-STIM; *bottom*). Note the place-field stabilization for INT-STIM condition. (**C**) Population vector correlation across trials for control and INT-STIM conditions. (**D**) Population vector (PV) correlation was significantly higher for INT-STIM compared to control (\*\*\*\**p* = 3.71e-25, Wilcoxon paired test, *n* = 190 position bins). (**E**) Schematic of experimental design (see also Figure S6C). (**F**) Trial-by-trial rate maps of an example CA1 place cell. (**G**) Population vector correlation during PRE, PV-STIM, and POST trials for inbound (IN) and outbound (OUT) trajectories on the linear track in an example session (no stimulation during PRE and POST blocks). Note higher population vector correlation during PV-STIM blocks. (**H**) Population vector correlation during PV-STIM compared to PRE and POST (\*\*\*\**p* = 4.5e-10 and 6.08e-7, Wilcoxon paired test, *n* = 9 sessions from 4 mice). (**I**) Neural trajectories of left-bound trials during figure-8 alternation task (Figure 1C) for an example session. Hollow circles for correct trials, solid circles for error trials (trial 6 and 22). (**J**), Detailed view of one error trial shown in (**I**; trial 6). While trial 5 and 6 were closer in time compared to trial 6 and 8, a jump in neural distance was observed, even before the choice point (CP). (**K**) Euclidean distance of neural trajectories between correct trials and adjacent error or correct trials (*n* = 28 sessions from 6 mice, each paired dot line for one session, \*\*\*\**p* = 3.17e-9, 4.04e-7 for complete trajectory and segment before CP, respectively, Friedman test with Dunn’s *post hoc*). (**L**) *Left*, Schematic of the perturbation protocol. *Right*, Task performance difference between PV-STIM and control sessions (n.s., *p* = 0.53, *\*p* = 0.042, rank-sum test compared to 0; *\*p* = 0.049, rank-sum test compared WT vs. PV::ChR2; *n* = 26 sessions from 5 WT mice, and 27 sessions from 5 PV::ChR2 mice, respectively). WT, wild type. See also **Figure S6**.

To experimentally test the role of CA1 inhibition in place cell drift, we performed an optogenetic manipulation in mice equivalent to the one in the model. We focused on one specific subtype of interneurons, parvalbumin expressing -PV-interneurons, as previous work showed their role in modulating place cell spike timing ^66,76,79^. We trained transgenic mice expressing channelrhodopsin (ChR2) in PV interneurons to run back and forth on a linear track (Figures 5E and S6B). In a subset of trials, we activated PV interneurons with low-intensity, white noise-modulated, blue light to disrupt their endogenous spike timing (Figure 5E). Similar to the model manipulation, this optogenetic manipulation enhanced the moment-to-moment variability of inhibition and the correlation among PV interneurons (Figure S6C). The manipulation did not affect animal locomotion (Figure S6B), basic place field properties, or remapping across different trajectories and mazes (Figures 5H and S6C-S6H). However, place field stability significantly increased during the manipulation (Figure 5F-5H), suggesting a reduced population representational drift (Figure S6E), similar to our simulation results (Figures 5D and S6A). Furthermore, standard metrics of spatial coding, like spatial information, were enhanced during optogenetically stimulated trials (Figure S6D). Together, our simulations and experiments indicate that local PV interneurons actively promote representational drift in CA1 place cells, an effect that was dissociated from place field spatial remapping.

Overall, our results show that place cell spatial remapping and representational drift are driven by feedforward excitatory inputs and local inhibition, respectively. This dissociation provides mechanistic support for our hypothesis that population representations of space and time are disentangled in CA1.

Finally, we sought to further establish the relationship between CA1 disentangled population dynamics and behavioral performance. Given that the manipulation of PV cells provided a way to specifically disrupt representational drift while preserving place cell tuning, we focused on this feature of CA1 dynamics. First, we compared correct and error trials in the original delayed alternation task. Neural trajectories for error trials displayed an abrupt jump on the manifold respect to adjacent trials, deviating from the smooth temporal progression of correct trials (Figures 5I-5J and S6I). To quantify this observation, we calculated the distance of neural trajectories between adjacent trials and found that it significantly increased during errors (Figure 5K). Importantly, this deviation was already present in the central stem of the maze, before reaching the choice point (Figures 5J-5K), thus preceding an incorrect choice. This jump was not caused by differences in running speed (Figure S6J), nor the neural trajectories for error trials were off-manifold (Figures 5I and S6K). This result suggested a relationship between the temporal structure of representational drift and successful memory-guided behavior in this task. Therefore, we sought to prove this relationship by delivering, in a random subset of trials, the same noise stimulation to PV interneurons as before (Figure 5E) while animals performed the spatial alternation task (Figure 5L). This perturbation increased the fraction of errors compared to non-stimulated trials (Figure 5L). The same light stimulation in control mice (not expressing ChR2) had no effect on performance (Figure 5L), suggesting that CA1 representational drift contributes to memory-guided navigation.

## Discussion

The ability to rapidly generalize common elements across experiences and use this knowledge to flexibly guide new behaviors is a fundamental cognitive ability of humans and other animals. Using large-scale neural ensemble recordings and manipulations in mice performing a similar task across different mazes, we identified the mechanisms that enable the hippocampus to rapidly generalize shared features across experiences while also preserving their individual specificity. Population dynamics disentangled task variables (location, time and choice), and these representations generalized across environments, whereas individual neurons responded conjunctively (i.e., with mixed selectivity) to different features on each maze. Therefore, memory specificity and rapid generalization are supported by disentangled population dynamics with conjunctive single-cell responses in the hippocampus.

Extensive prior work, largely conducted in rodents, established that hippocampal representations are characterized by two main features that support their role in encoding and discriminating episodic memories. First, each experience is encoded by the activity of a sparse subset of cells in a conjunctive manner, combining different aspects of an experience, such as location, time and content, into a unified representation ^2,10,26-28,30,35,37,44^. Second, ensemble patterns for different experiences tend to be maximally uncorrelated (i.e., pattern separation) ^4,31-33,35,37^. Such representational properties make it challenging to generalize across experiences, as their individual representations are statistically independent. On the other hand, recent work, mainly in humans and other primates in non-spatial task settings, has shown that the hippocampus generates abstract representations of tasks variables that support inference, generalization and concept learning ^7,8,22,23,80-82^. Several possible explanations may account for these unresolved discrepancies, including possible differences in hippocampal functions across species, inference and episodic memory requiring different cognitive demands supported by distinct hippocampal representations, or gradually formed abstract representations in the neocortex that subsequently feedback into the hippocampus.

Our results support an alternative hypothesis that can unify prior findings in rodent work on episodic memory and spatial navigation with primate work on abstraction and inference, while also explaining the role of the hippocampus in rapid generalization. We found generalized coding at the population level of task variables across different mazes, which enabled rapid generalization and efficient learning, despite the seemingly random reorganization of single neuron responses between environments. Importantly, the geometry of the hippocampal neural manifold evolved with learning, reflecting mice’s ability to learn faster new tasks with the same latent structure as previous ones. This transfer learning ability could be supported by the generalization of low-dimensional representations of task structure across different mazes ^28,42^. We thus propose that a disentangled, generalized population code is a fundamental property of the hippocampus across cognitive demands and species. In turn, these hippocampal representations are broadcasted to the neocortex where they can serve as the basis for the gradual development of schemas, abstract concepts and categories ^7,9-11,19,54,59,83^.

Two critical challenges for maintaining generalized representations across environments, or even a stable representation of the same environment over time, are the seemingly random and uncorrelated nature of spatial remapping ^33,34,38^ and temporal drift ^38,45,84^. In contrast to this notion of complete random hippocampal remapping and drift, we found that both processes are highly structured at the population level. We found that they are supported by distinct circuit mechanisms: synchronous excitatory inputs and precisely timed local inhibition, respectively. It remains to be elucidated how these mechanisms interact with other single cell and circuit processes shown to also influence place cell remapping and drift ^65,67,85-87^. In particular, we found that assemblies of coactive CA3-CA1 cells remapped in a coordinated manner across mazes. This correlated remapping could underlie the preserved manifold structure and coherent coding dimensions observed across environments, in line with recent proposals ^28,64^.

In summary, our study advances a novel framework for understanding the structure of hippocampal episodic memory representations and how they can support both discrimination and rapid generalization, key elements for intelligent behavior. The network mechanisms and dynamics uncovered in this work can inform the design of artificial neural networks that currently struggle to achieve rapid generalization efficiently and to transfer acquired knowledge to solve new problems.

## Acknowledgments

The authors thank members of the Oliva and Fernandez-Ruiz labs, Jesse Goldberg, Weinan Sun and Daniel Levenstein for providing useful feedback on the manuscript, and Ipshita Zutshi for generously sharing data. This work was supported by NIH grant R00MH122582 and R01MH130367 (AO), NIH grant R01MH136355, DP2MH136496, Sloan Fellowship, Whitehall Research Grant, Klingenstein-Simons Fellowship and Pershing Square Foundation’s MIND Prize (AFR), and Klarman Fellowship (WT).

## Author contributions

W.T., H.C., A.O. and A.F.-R. conceived and designed experiments and analyses, analyzed data and wrote the manuscript. W.T., H.C., C.L., S.P.-H, W.Y.Z. and J.P. performed experiments. These authors jointly supervised this work: A.F.-R, A.O.

## METHODS

### Experimental model details

All experiments conformed to guidelines established by the National Institutes of Health and have been approved by Cornell University’s Institutional Animal Care and Use Committee. All mice were kept in the vivarium on a 12-hour light / dark cycle with *ad libitum* access to food and water except during training. Temperature and humidity in the room were kept at 68-72 F and 40-60%, respectively. They were housed with a maximum of 5 per cage before surgery and individually afterward. C57BL/6J and B6FVBF1/J male mice (∼27-32 g, 3-6 months old, The Jackson Laboratory) were used for electrophysiological recordings, and Pvalb-IRES-Cre/+::Ai32/+ mice (∼26-30 g, 3-8 months old) and C57BL/6J (∼26-30 g, 3-8 months old) wild type mice were used for optogenetic manipulation and control experiments. The former line was generated by crossing B6.129P2-Pvalbtm1(cre)Arbr/J and Ai32(RCL-ChR2(H134R)/EYFP (The Jackson Laboratory). Tg(Grik4-cre)G32-4Stl/J (The Jackson Laboratory) mice were used for the CA3 chemogenetic experiments.

### Surgical procedures

Silicon probes were implanted as previously described ^1,88,89^. Briefly, mice were anesthetized with isoflurane, coordinates were taken following stereotaxic guidance after which craniotomies were performed. Silicon probes (NeuroNexus or Diagnostic Biochips) were mounted on micro-drives to allow accurate adjustment of the vertical position of the electrodes after implantation. A variety of electrodes were used for these experiments, including A5×12/16-buz/lin-5mm-100-200-160 (64 channels), ASSY-64-2-6 (128 channels), and A4×32-Poly2-5mm-23s-200-177 (128 channels). A stainless-steel wire connected to a thin wire (California wires) was inserted over the cerebellum and used as both ground and reference and cemented in place using C&B Metabond (Parkell). A layer of Metabond was applied to the full extent of the skull and a customized 3D-printed plastic head-stage was attached to the skull using Metabond and dental acrylic, over which four flaps of copper mesh were attached to cover the implant. The probe(s) was then inserted right above dorsal CA1 (AP = -1.9 mm, ML = 1.6 mm from Bregma and 0.9 mm from the surface of the brain); aided by the micromanipulators. For CA3-CA1 dual recordings, one A2×64-Poly2 electrode was inserted to target both dorsal CA1 and CA3 (−2.0 mm AP, +2.0 mm ML and 1.0 mm from the surface of the brain) in one hemisphere and another electrode in the opposite hemisphere to target either CA1 or also both CA1 and CA3. Once the probe had been inserted and drives cemented to the skull, craniotomies were sealed with artificial dura-gel. For animals used for optogenetic manipulations (see *optogenetics and chemogenetic manipulations*), an optic fiber (200 μm core) was attached under microscope guidance to one of the shanks of the silicon probe and implanted in CA1 (AP = -1.9 mm, ML = 1.6 mm) one hemisphere. In the contralateral hemisphere, a second optic fiber was implanted over the CA1 and cemented to the skull. In a different set of animals, optic fibers were bilaterally implanted over medial entorhinal cortex (AP = +0.2 mm, ML = ± 3.75 mm and 0.6 mm from the surface of the brain, with angled 6 toward anterior) ^73^. Finally, the four flaps of copper mesh were bent upwards and soldered between them to provide stability to the implant. The ground wire from the cerebellum was connected to the ground and reference wires of the probes and all together to the copper mesh. Post-operative care included administration of carprofen subcutaneously for 3 days and triple antibiotic ointment. Mice were allowed to fully recover in their home-cage over a heating pad. After recovery (∼1 week), the probe was lowered gradually in 75–150-μm steps per day until the desired position was reached. We used physiological landmarks and characteristic LFP patterns to identify the layers corresponding to the different hippocampal subregions ^1,66,88,90,91^.

### Behavioral tasks

Before surgery, animals were accustomed to one maze for 3-5 days. After surgery recovery, mice were handled daily and accommodated to the experimenter, recording room, and cables for at least a week before the start of the experiments. One day before the beginning of the training, animals were placed under water restriction.

In the delayed spatial alternation task (Figures 1C and 2A), mice (*n* = 6) were trained to run in the figure-8 maze, starting at the beginning of the center stem and having to alternate between left and right arms of the maze in each trial. An 8-s delay was introduced at the beginning of each trial, during which the animal was confined at the center stem by automatic doors. Upon completion of a correct trial, a small reward (30% sugar water) was provided at the end of the side arms. Maze doors and water delivery were fully automated with a custom circuit. Animals ran two or three behavioral sessions per day (∼20-40 trials for ∼30 mins per session), interleaved with 2-hour sleep sessions.

In the generalization task, mice (*n* = 5 for M1 and M2, *n* = 4 for M1, M2 and M3) were first trained on the delayed alternation task on one maze (M1, 61-cm diameter), as described above. After mice consistently surpassed 80% of correct trials per session (final performance = 83.9 ± 3.2%, as mean ± sem for 5 mice), which took 4-5 days, the second figure-8 maze (M2) was introduced. These two mazes had different spatial layouts (Figure 1C), local cues (textures, walls and landmarks) and were located in different rig enclosures with distinct distal cues. On each day, mice ran in M1 and M2 in morning and afternoon sessions (in a variable order) separated by a 2-hour rest period in their home cage. After 2-3 days of the M1-M2 task, a third maze (M3, square shape, 40-cm diameter, Figure 2A) was introduced. M3 was located in a different room. For the 3-maze generalization test, each day, mice were trained on the same alternation task in each of these three different mazes for one session each (∼25-30 trials per session) in random order of presentation, with 2-hour intervals between sessions.

In the forced alternation version of the task only one of the two side arms remained open in each trial. The open arm switched to the opposite one in each trial (without delay). Therefore, animals (*n* = 5) were forced to alternate between left and right arms but, contrary to the standard version of the alternation task, in this case, there were no memory demands. Mice obtained a water reward at the end of each side arm.

In the linear-track task (Figure 5E), mice were trained to run back and forth to collect sugar water rewards at both ends of a 1m-long track. Rewards were automatically delivered in reward wells. Animals (*n* = 4) ran two sessions per day (∼30 min and ∼30 trials per session), interleaved with 2-h sleep sessions. After running for at least one week in the first linear track, a second linear track (1m-long) located in a different room was introduced (Figures S6G and S6H). On some days, mice ran morning and afternoon sessions in each of these two tracks in a random order.

In all the behavioral sessions, animals’ position was tracked with an overhead camera (Basler acA720-520uc and acA1300-200uc) at 30 Hz (Pylon 5.0 software), and later extracted using DeepLabCut (DLC, v2.3.0) ^92^.

### Optogenetic and chemogenetic manipulations

In PV::ChR2 animals, white noise light pulses were delivered using a 470-nm LED (Prizmatrix) in a subset of trials while they were running in either a linear track (*n* = 4 mice) or figure-8 maze (Figure 5; *n* = 5 mice). A patchcord was connected to a commutator (Prizmatrix) on top of the maze. Light intensity was low (maximum intensity of 3 mW), but it visibly entrained interneuron spiking in CA1. On the linear track, no stimulation was given during the first third and last third of trials (PRE and POST trial blocks, Figure 5E), serving as control conditions. Stimulation was only delivered during the middle third of trials (PV-STIM block) and exclusively during running periods. In the figure-8 alternation task (Figure 5L), the same optogenetic stimulation was delivered in a random selection of 3 out of 6 trial blocks (approximately 5 trials per block). The remaining 3 blocks were without stimulation (i.e., control trials). This protocol was also applied to wild-type (WT) mice (*n* = 5), similarly, implanted with optic fibers but not expressing ChR2, to control for nonspecific effects of the optogenetic stimulation.

To test the impact of excitatory inputs on CA1 representational drift and remapping, we re-analyzed the dataset previously published in Zutshi *et al*. ^73^ with bilateral CA3 chemogenetic (Figure 4) and bilateral MEC optogenetic (Figures S5C-S5G) manipulations. In brief, mice were implanted with 64 or 128 channel silicon probes in dorsal CA1 and trained to run on the delayed spatial alternation task in a figure-8 maze (Figure 4E; for more details of the procedures, see Zutshi *et al*. ^73^). All the mice were trained to obtain a minimum performance of 80% before the start of the experiments. During the experiment days, animals ran 2 sessions per day, interleaved with 3-4 hour rest in the home cage. For the CA3 disruption cohort of mice, 4 Tg(Grik4-cre)G32-4Stl/J mice were injected with AAV5-Syn-DIO-PSAM4-GlyR-IRES-GFP (titer: 2×10^12^, gift from Scott Sternson, Addgene viral prep #119741-AAV5; RRID:Addgene_119741) bilaterally into CA3, to express the chemogenetic inhibitor PSAM4 ^74^. After the first behavioral session, mice were given an i.p. injection mid-way through the home cage rest session, with either 0.9% saline (Lifeshield Diluent NaCl), or the PSAM4 activator μPSEM792 hydrochloride (Tocris, #6865), 1.5-3 mg/kg diluted in 0.9% saline (working concentration 1 mg/ml, volume injected: 0.04-0.06 ml). For MEC disruption (Figures S5C-S5G), 3 mice were bilaterally injected with AAV5-mDlx-ChR2-mCherry in MEC. MEC disruption was achieved by implanting 200-μm optical fibers bilaterally over the MEC. Light stimulation was calibrated before the experiment (2 mW to 8 mW) and was delivered in blocks of 10 trials interleaved with 10-trial blocks of no stimulation, restricted to the central arm of the maze.

### Recording system and data preprocessing

An Intan RHD2000 interface board or Intan Recording Controller was used for electrophysiological recordings. The sampling rate was set at 20 kHz. Both amplification and digitization were done in the head stage (Intan Technologies). Data were visualized online during recording using the Intan software and Neuroscope (Neurosuite). LFP signals were down-sampled at 1250 Hz for subsequent analysis.

### Tissue processing and immunohistochemistry

After experiments were terminated, mice were deeply anesthetized and kept under high-flow (4-5%) isoflurane anesthesia. They were then perfused transcardially with 0.9% phosphate buffer saline (PBS) solution followed by 4% paraformaldehyde (PFA) solution. Brains were kept in PFA for 24 hours and if necessary, in PBS afterwards until further processing. Brains were then sectioned into 70-μm thick slices (Leica Vibratome, 2000). Finally, sections were washed and mounted on glass slides with fluorescence medium (Fluoroshield with DAPI – F6057, Sigma, USA). Immunostained slices were examined, and histological images were acquired with a Zeiss confocal microscope.

## Quantification and statistical analyses

### Spike sorting and single unit classification

Spike sorting was performed semi-automatically using KiloSort ^93^ (https://github.com/cortex-lab/KiloSort), followed by manual curation using the software Phy (https://github.com/kwikteam/phy) and custom designed plugins (https://github.com/petersenpeter/phy-plugins) to obtain well-isolated single units. Cluster quality was assessed by manual inspection of waveforms and auto-correlograms, and by isolation distance metrics. Multi-units, noise clusters, or poorly isolated units were discarded from further analysis. Well isolated units were classified into putative cell types using the MATLAB package, Cell Explorer ^94^ (https://github.com/petersenpeter/CellExplorer). Spiking features such as auto-correlogram (ACG), spike waveform, and putative monosynaptic connections derived from short-term cross-correlograms (CCGs), were used to characterize and classify well-isolated units ^95^. Three cell types were assigned: putative pyramidal cells, narrow waveform interneurons, and wide waveform interneurons. The two key metrics used for this separation were burst index and trough-to-peak latency. Burst index was determined by calculating the average number of spikes in the 3-5 ms bins of the spike ACG divided by the average number of spikes in the 200-300 ms bins. To calculate the trough-to-peak latency, the average waveforms were taken from the recording site with the maximum amplitude for the averaged waveforms of a given unit. Only putative pyramidal cells were used for further analysis, unless otherwise specified.

### Spatial fields and linearization

Spatial fields were calculated only during running periods (> 2 cm/s) at positions with sufficient occupancy (> 20 ms). 2D occupancy-normalized rate maps were constructed using spike counts and occupancies with 150 × 150 spatial bins, smoothed with a 2D Gaussian kernel (SD = 2). To construct the 1D linearized rate maps on the 2 trajectory types in a figure-8 maze (left vs. right), animals’ linear positions were first estimated by projecting the actual 2D positions onto pre-defined idealized paths along the track, and further classified as belonging to 1 of the 2 trajectory types ^1^. The linearized rate maps were then calculated with 100 equally spaced bins of the linear positions and smoothed with a 1D Gaussian (SD = 1).

### Linear-nonlinear (LN) model

To quantify the relationship between single-neuron spiking and different behavioral variables (position, trial time, and behavioral choice, or P, T, and C), we used a LN model framework (Figure S1) ^96^. The LN models estimate the spike rate of a neuron at time bin *t* (*r*_*t*_) as an exponential function of the sum of all variables (*P*_*t*_, *T*_*t*_, *C*_*t*_) projected onto a corresponding set of parameters (*β*_*P*_, *β*_*T*_, *β*_*C*_) as:

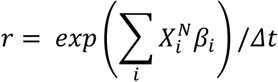

where ***r*** denotes a vector of firing rates of a neuron across *N* time bins, *i* indexes the variables (*i* ∈ [*P,T,C*]), and 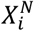 is a state matrix for the *i*-th variable. Each column of 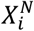 is a state vector *x*_*i*_ at each of the *N* time bins, whose elements are 0, except for the element (= 1) corresponding to the animal’ current state of that variable.

To learn the parameters (*β*_*P*_, *β*_*T*_, *β*_*C*_), we maximized the Poisson log-likelihood of the observed spike train given the model estimated spike number 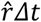. The parameters were optimized using *fminunc* function from MATLAB, and the log-likelihoods were calculated from the held-out data via 10-fold cross-validation. To select the best LN model that characterizes the single-neuron response (Figure S1D), we considered 3 single-variable models (*P, T, C*), 3 double-variable models (*PT, TC, PC*), and 1 full model (*PCT*), and selected the best one as previously reported ^96^. In brief, we first determined which single-variable model had the highest performance (i.e., highest log-likelihood), and then compared this model with all the double-variable models. If one of the double-variable models performed significantly better than the best single-variable model, we further compared the double-variable model with the full model. In all cases, significance was quantified through a one-sided rank sum test with *p* < 0.05 ^96^.

### Choice selectivity

To measure choice selectivity of each neuron (Figure S1H), we calculated the mean firing rate of the neuron on the center stem before the choice point (*FR*) when animals ran on the L versus R trajectory as:

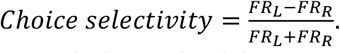

The correlation of single-neuron choice selectivity across two different mazes were calculated (Figure S1H). The significance of single-neuron choice selectivity was further assessed by shuffling the choice labels across different trials; a cell with *p* < 0.05 was defined as a choice-selective cell (or splitter cell; Figure S3A) ^97-99^.

### Population vector correlation

To measure remapping in population activity (Figures 5, S1 and S4-S6), a population vector (PV) was constructed as the firing rate vector of all spatial-tuned cells in a certain linear spatial bin. The PV correlation was then defined as the Pearson correlation (*r*) between the PVs across all bins in the mazes between two sessions, or across two different trials.

### Representational dissimilarity matrix

To assess the similarity between trial time and neural representational drift (Figures 1H and 1I), we created a representational dissimilarity matrix (RDM) that consisted of the Euclidean distance between the population vectors or single-unit firing rate vectors for different trials (Figure 1H, *right*). Similarly, we created a RDM for trial time using the median timestamp of each trial and calculating the time difference between each pair of trials. The upper triangular parts (without the diagonal) of the neural RDM and the RDM of trial time were extracted to compute a Spearman correlation coefficient between them (Figure 1I).

### Visualization of neural manifolds

All the simultaneously recorded cells that pass the cell inclusion criteria (see *spike sorting and single unit classification*) were included in manifold analyses. For each behavioral session, spike counts were taken in 200-ms bins and smoothed with a 1-s moving window (*locsmooth* from the Chronux toolbox, http://chronux.org/, V2.12). Only running periods (running speed > 2 cm/s) were included, and bins with spikes from less than 5 cells were discarded. Uniform Manifold Approximation and Projection (UMAP; https://www.mathworks.com/matlabcentral/fileexchange/71902-uniform-manifoldapproximation-and-projection-umap, V4.2.1) ^100^ was then run on these *N*-dimensional data (*N* for number of simultaneously recorded cells) without any labels (i.e., unsupervised) to extract low dimensional neural manifolds (e.g., Figures 1F and 1G). The hyperparameters for UMAP were: ‘n_dim’ = 3, ‘metric’ = ‘cosine’, ‘n_neighbours’ = 40, and ‘min_dist’ = 0.6, similar to previous studies ^11,61^.

### Principal angles between subspaces

To characterize how different coding subspaces were oriented relative to each other in neural state space, we computed their principal angles (Figure 1J) ^55,57,58^. Specifically, we first constructed a population activity matrix for each of the three different task features (i.e., location, time and choice type). The activity matrices for location and time were constructed using the trial-averaged and location-averaged linearized rate maps of all cells, respectively, whereas the choice matrix was from the firing rate difference between left and right trials on the center stem (i.e., splitterness) ^97-99^. For a given activity matrix (**X**), we used singular value decomposition (SVD) to find a rank-2 approximation as **X** = **U**\***S**\***V**. Then, for two given features *a* and *b* with the associated 2-dimensional bases 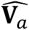 and 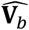, a SVD was performed onto their inner product matrix 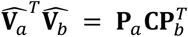, and the first principal angle was reported as:

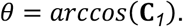

### Alignment between coding directions

We examine how much the coding direction of one variable aligns with the direction of another variable (Figures S2B-S2E) we used a previously described method ^101^. In brief, let *z(x)* denote the neural population activity, where *x* reflects the maze position. The changing direction across time (i.e., drift direction; *Δμ*_*z(x)*_) was defined as the change in the mean neural activity *μ*_*z(x)*_, and the changing direction across positions (i.e., spatial coding direction), was defined as ∇_*x*_*z(x) =* Δ*z*(*x*)/Δ*x*. Similarly, the changing direction across choice (i.e., splitting direction) was defined as the difference in neural population activity between the L and R trajectory before the choice point *z*_*L*_(*x*) -*z*_*R*_(*x*), where *x* < choice point. We then computed the root-mean squared dot product between two coding directions. For example, for the drift and spatial coding directions, the root-mean squared dot product was computed as:

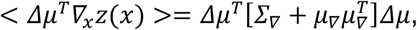

and then the dot product was normalized by its largest eigenvalue *λ*_*max*_ as:

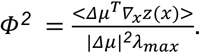

Finally, the alignment coefficient *ρ* was computed by comparing *Φ* with the *Φ*_*0*_ expected from two randomly oriented vectors as 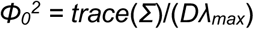 as:

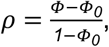

such that *ρ* = 1 is for perfect alignment and *ρ* = 0 is for randomly oriented vectors.

### Gain modulation approximation

As an alternative measure of the subspace overlap, we asked how well a simple gain-modulation model can approximate the full linear regression model at the population level, as previously described ^55^. In brief, we approximated population neural activity ***κ***(*t,p*) with a gain modulation formula:

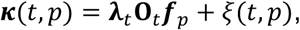

where **λ**_*t*_ is the modulation factor for the trial *t*, ***f***_*p*_ is the spatial location vector independent of trial numbers, **O**_*t*_ is the orthogonal matrix, and *ξ* (*t, p*) is the approximation error. For a given trial *t*, the similarity score between ***κ***(*t*,:) and the approximation is defined as:

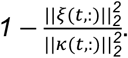

### Multidimensional scaling

Embeddings of population activity via multidimensional scaling (MDS; Figure 2) were computed as follows ^62^. First, we computed the trial-by-trial rate maps for all cells. Next, we calculated the rate map correlation across cells for every pair of trials and transformed this correlation matrix into a distance matrix by computing one minus this correlation matrix, with the diagonal = 0. Finally, this distance matrix was reduced to a 2-dimensional embedding via MDS with the metric stress criterion (*mdscale* from MATLAB). The left and right maze arm trials were embedded separately and simultaneously for trial time (Figure 2B) and choice types (Figure 2C), respectively. To calculate the angular difference between the embeddings for two different sessions (Figure 2H), the average changing direction *θ* for a given session was calculated across all the trials as:

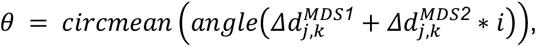

where *i* is the imaginary unit, and 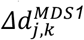 and 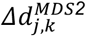 are the changes in MDS1 and MDS2 between the *j*-th and *k*-th trial. The angular difference in the changing directions between the session 1 and 2 was then calculated as *Δθ* = *abs*(*θ*_*2*_ − *θ*_*1*_) using *angdiff* from MATLAB.

### Cross-condition generalization performance

The property of neural manifolds to generalize to unseen experimental conditions was estimated using cross-condition generalization performance (CCGP) ^7,8,11^. For choice types (L vs. R), we first trained a binary linear decoder to classify choices using data from one maze and tested the trained decoder on data from the other two mazes that were not used for training (Figure S3E). A binary decoder was built using a *C*-Support Vector Machine (SVM) with a linear kernel with a 4-fold cross-validation through the *libsvm* library (https://www.csie.ntu.edu.tw/~cjlin/libsvm/, V3.12) ^11,102^. Given that trial time is a continuous rather than discrete variable, we used a regressor based on the first principal dimension (PC1) from the M1 data, which captured the most variance across M1 trials, and tested the generalizability by projecting the neural activity from M2 or M3 onto the M1 PC1 (*linproj* from MATLAB; Figure S3D). Fitting accuracy was then assessed by calculating the Spearman correlation coefficient between the length of the projection and the actual trial time. For both time and choice decoding, the actual CCGP was tested against its condition-label permutations (*n* = 1000 times). Only sessions with at least 10 trials for each choice type and with sufficient accuracy on the training dataset (decoding accuracy ≥ 70% and *p*-value < 0.01 for the correlation coefficient for choice and time, respectively) were included.

### Neural manifold alignment

To answer the question of whether the geometry of the latent task structure was preserved across different mazes, we aligned the low-dimensional neural manifolds from two mazes using a Procrustes transformation (Figures 2J, 2K and S3F; *Procrustes* from MATLAB) ^63^. A Procrustes transformation involves translation, scaling, and rotation, and thus preserves the geometry of the neural manifold. Specifically, we calculated the centroids of the neural activity (in 200-ms bins) for each position bin in a maze (*n* = 100 position bins), and this provided an estimate of the manifold geometry as a 100-point cloud for each maze embedded in a 3-dimensional space. A Procrustes transformation was applied to find the optimal translation, scaling, and rotation matrices that align these two point clouds. The moment-by-moment neuronal population activity (in 200-ms bins) from a different session (M1’, M2 or M3) was then aligned to the maze 1 (M1) activity using the optimal transformation matrix (Figure S3F). After alignment, we used a *K*-Nearest Neighbor (KNN) classifier (*K* = 100 position bins) built from M1 data to decode positions of the aligned M1’, M2 or M3 data (*fitcecoc* with KNN Learners and *predict* from MATLAB; Figures 2J, 2K and S3G). The performance of the decoder was defined as the correlation coefficient between the actual position of the animal and the location predicted by the decoder.

### Bayesian decoding

The decoding based on manifold alignment above was compared with the canonical Bayesian decoding of animal location in each maze ^1,103,104^. For each time bin (bin = 500 ms; 200-ms bins yield similar results), a memoryless Bayesian decoder was built for the L- and R-trajectory types to estimate the probability of animals’ position given the observed spikes (Bayesian reconstruction; or posterior probability matrix):

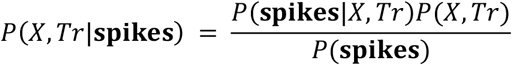

where *X* is the set of all linear positions on the track for the 2 trajectory types (i.e., *Tr*, L or R), and we assumed a uniform prior probability over *X* and *Tr*. Assuming that all *N* cells active in a sequence fired independently and followed a Poisson process:

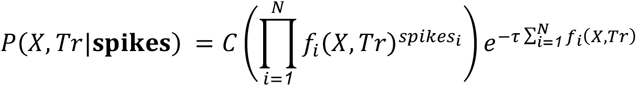

where *τ* is the duration of the time window (*τ* = 500 ms), and *C* is a normalization constant such that 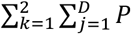 *P*(*x*_*j*_, *tr*_*k*_ **| spikes)** = 1 (*x*_*j*_ is the *j*-th position bin, *D* is the total length of the track, and *tr*_*k*_ is the *k*-th trajectory type; *k* = 1 or 2, representing L or R trajectory, respectively). The Bayesian decoder was built based on the firing rate maps, *f*(*X,Tr*), in M1, and was then applied to the *spikes* from M1’ or M2 (Figure 2J). Consistent with the alignment procedure, the performance of the decoder was defined as the correlation coefficient between the actual position of the animal and the location predicted by the decoder.

### Persistent (co)homology

Persistent (co)homology is a tool in topological data analysis with recent success in describing the ring topology of head direction cells ^60^, torus of grid cells ^61^ and place fields in the hippocampus and subiculum ^49^. In brief, we calculated the centroids of the neural activity for each position bin in a maze (*n* = 100 bins) embedded in the 2-dimensional UMAP space, which provided an estimate of the whole manifold geometry (see also *neural manifold alignment*). This downsampling is also necessary for the robustness of the co-homological analysis ^105^. Then, each point in the 100-point cloud was replaced by a ball of infinitesimal radius, which was gradually expanded at unison. Taking the union of balls of a given radius results in a space with holes (or loops) of different sizes. The range of radii for which the loop is born (or appear) and dead (or disappear) is referred to as the “lifespan” of the loop (Figure 2C). The MATLAB toolbox *PersistentHomologyOnMATLAB* (https://www.mathworks.com/matlabcentral/fileexchange/129579-persistent-homology-based-on-alpha-complex-on-matlab) was used to calculate persistent (co)homology. Finally, the lifespan index (Figures 2D and 2E) was calculated as the lifespan ratio between the first and second most prominent loops of the first cocycle (*H1*).

### Hippocampal network models

We adapted linear Hebbian/ anti-Hebbian network models developed for modeling representational drift in a previous study ^70^. In brief (for a more detailed description, see Qin *et al*. ^70^), this network minimizes the mismatch between the similarity of pairs of inputs (***x***) and corresponding pairs of outputs (***y***, representing neuronal firing rate):

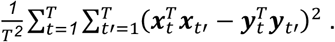

Notably, the cost function above is rotational symmetry. That is, for inputs with the same geometric structure (i.e., two circular mazes), if a set of outputs {***y***^1^, …, ***y***_*T*_} is an optimal solution to this problem, {**R*y***_1_, …, **R *y***_*T*_} will also be an optimal solution (where **R** is a rotational orthogonal matrix), which is consistent with the rotational transformation of neural manifolds across mazes observed in the experimental data (Figures 1F and 1G). The optimization of this cost function is learned by the network, where each output *y*_*t*_ is produced by running the following neural dynamics until convergence:

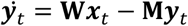

where **W** and **M** hold the synaptic weights of feedforward and lateral connections, respectively. After the convergence of the neural dynamics, **W** and **M** are updated with a Hebbian and an anti-Hebbian rule, respectively, with synaptic noise (*ξ*):

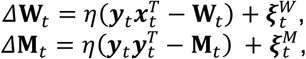

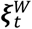 and 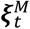 are independent Gaussian noises. To further explore how different sources of synaptic plasticity in the network contribute to remapping (rotational dynamics of neural manifolds; Figure 4) and temporal drift (parallel shift of neural manifolds; Figure 5), we manipulated feedforward or local noise (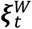 or 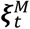) after the convergence. For the manipulation to the local inhibition, we injected a shared white noise pattern to all lateral synapses, resulting in a decrease in 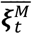, similar to our experimental perturbation of local PV neurons. For the input manipulation, we inhibited feedforward input **W**, and amplified the forward noise (i.e, 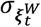), in a similar way to the experimental manipulations of CA3 in Figure 4.

For the CA3-CA1 network model (Figure 3), we constructed two layers of neurons as CA3 and CA1, where the CA3 layer (***y***) was updated based on spatial inputs as a circular maze (***x***) as 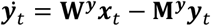, and the CA1 layer (***z***) was updated based on CA3 layer inputs as 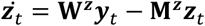 We reduced **M** of the CA3 layer (**M**^***z***^ = 0.1**M**^***y***^) to reflect the strong recurrent excitatory connections within CA3.

### Cross-regional cell assembly detection

The cross-regional cell assembly detection method was implemented as previously described ^104,106^. This method is equivalent to Principal Component Analysis (PCA) based within-regional cell assembly detection ^107^ but uses Singular Value Decomposition (SVD) to diagonalize the non-square cross-regional correlation matrix. In brief, spike trains of each neuron during running periods (running speed > 2 cm/s) were binned into 50-ms intervals for the whole session and then *z*-transformed as 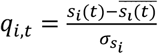, where *s*_*i*_(*t*) is the binned spike train of the *i*-th cell, and 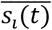 and 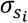 are the mean and standard deviation of *s*_*i*_(*t*). For simultaneously recorded *N* CA1 and *M* CA3 cells, a (*N* x *M*) correlation matrix (**C**) was calculated for all CA3-CA1 cell pairs, with each element (*c*_*i,j*_) representing the Pearson correlation coefficient of the *i*-th CA1 cell and the *j*-th CA3 cell as *c*_*i,j*_ *= q*_*i*_^CA1^*q*_*j*_^CA3^/*T* (*T* is total number of time bins). Then, SVD decomposes the (*N* x *M*) correlation matrix (**C**) into **U**\***S**\***V**, where **U** (*N* x *N*) and **V** (*M* x *M*) are the eigenvectors of **CC**^*T*^ and **C**^*T*^**C**, respectively, and **S** is an (*N* x *M*) matrix containing the singular values (analogous to eigenvalues) along its diagonal. Therefore, the columns of **U** and **V** contain the “neuronal weights” (or loadings in the context of PCA) of CA1 and CA3 cells, respectively. When defined in this way, the “neuronal weights” form an orthogonal set of vectors (i.e., orthogonal assemblies). To detect meaningful assemblies, we only considered the significant components with relative variance larger than 0.7/*n* (*n* is the total number of components) ^108^ Most of the detected assembly patterns consisted of a few neurons with high weights and a large group of neurons with weights around zero (Figures 3G, S4D and S4E). We thus detected assembly members as cells with weights exceeding the mean weights across all cells by 4 standard deviations, independently for each detected assembly ^1,103^.

### Place-field center-of-mass shift

To compare place-field remapping across mazes for assembly members and non-members (see *cross-regional cell assembly detection*; Figures 3D and 3I), we calculated the center-of-mass (COM) of place fields for each maze as ^103^:

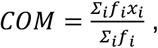

where *f*_*i*_ is the firing rate in the *i*-th spatial bin, and *x*_*i*_ is the distance of the *i*-th spatial bin from the start point of a trajectory. COM shift was then measured as the difference between the first and second mazes (Figures 3E and 3J).

### Theta cross-covariance

As a complementary method to the place-field COM shift above, we also calculated the theta cross-covariance of cell pairs across mazes (Figures S4F and S4G). Theta periods were detected after filtering based on a speed criterion of > 2 cm/s. Standardized cross-covariance during theta periods was then computed for pairs of CA1 neurons as in previous reports ^104,109,110^. In brief, cross-correlation of spike trains was first computed using 10-ms bins, and cross-covariance was then estimated by removing the expected firing rate of coincidence in each bin, normalized by the mean firing rates of the neurons, the bin size, and the total length of the theta periods, and followed by smoothing (50-ms Gaussian window). The peak of the standardized theta cross-covariance was determined in a ± 200-ms window around the 0-ms lag, and the time offset (*τ*) of the peak was determined. Of note, the time offset of theta-covariance between two place cells is strongly correlated with the distance between the peaks of the two place fields ^104,111,112^. Thus, the difference between theta-covariance time offsets across two mazes serves as a complementary method to the place-field COM shift for assessing remapping.

### Communication subspaces

To capture the functional connectivity between CA3 and CA1, we identified their communication subspaces using Canonical Correlation Analysis (CCA) ^57,71^ and Reduced Rank Regression (RRR) ^58,72^. To measure the population correlation between CA3 and CA1, we used CCA (https://github.com/joao-semedo/canonical-correlation-maps) to find pairs of dimensions, one in each area, such that the correlation between the projected activity onto these dimensions is maximally correlated:

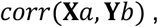

where **X** is a (*T* x *M*) matrix and **Y** is a (*T* x *N*) matrix containing the z-scored activity in the CA3 and CA1 population (as *q* in the *Cross-regional cell assembly detection*), respectively, with *T* representing the number of data points, and *M* and *N* representing the number of recorded neurons in each of the two areas, respectively. The vectors *a* and *b* have dimensions (*M* x 1) and (*N* x 1), respectively defining the dimensions in the population activity space of each area (i.e., communication subspaces). To quantify the stability of the communication subspaces across mazes, we tested whether the subspace (*a*_*M1*_, *b*_*M1*_) fit on one maze (M1) could generalize to another maze (M2). Specifically, we tested the cross-generalization of CCA:

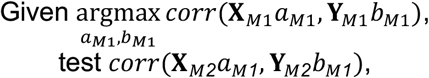

where **X**_*M*1_, **X**_*M*2_ and **Y**_*M*1_, **Y**_*M*2_ are the z-scored CA3 and CA1 population activity in M1 and M2, respectively. To test the significance of this correlation, we constructed a shuffle dataset by eliminating the moment-by-moment coordination across the two areas but preserving their overall activity statistics. Specifically, we circularly shifted the timestamps of each cell’s spike train in the M1 session (denoted as **X**_*shuffled M1*_ and **Y**_*shuffled M1*_), redetected the subspace (*a*_*shuffled M1*_, *b*_*shuffled M1*_) using **X**_*shuffled M1*_ and **Y**_*shuffled M1*_, and tested *corr*(**X**_*M*2_*a*_*shuffled M1*_, **Y**_*M*2_*b*_*shuffled M1*_) (Figure 3N, *right column*). To control for the possibility that the correlation observed reflected the shared spatial layouts across mazes, we identified the subspaces by replacing **X**_*M1*_ with linearized locations **L**_*M1*_ as (Figure 3N, *left column*):

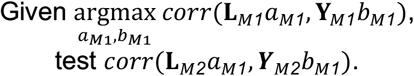

To investigate the dimensionality of the communication subspaces, we employed RRR as previously established ^72^ (https://github.com/joao-semedo/communication-subspace). In brief, similar to the CCA approach, we first related the moment-to-moment z-scored activity in the source CA3 population (**X**) to that in the target CA1 population (**Y**) using a linear model, **Y** = **XB**, where the coefficient matrix **B** is of size (*M* x *N*). RRR performed simultaneous dimensionality reduction and regression by imposing **B** to be a given rank *p*, and identifying a *p*-dimensional set of predictive subspaces in **X** that best predicts **Y**. This RRR can be solved using SVD as **B**_*RRR*_= **B**_*OLS*_ ***VV***^*T*^, where **B**_*OLS*_ is the least-squares solution and **V** is a (*N* x *p*) matrix with the top *p* columns representing principal components of the optimal linear predictor 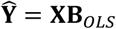. To find the optimal dimensionality for the RRR model (the value of *p*), we used 10-fold cross-validation and the smallest number of dimensions for which predictive performance was within one SE of the peak performance (Figure 3L) ^72^. As a complementary approach to CCA, we tested the cross-generalization of RRR ^58^ as 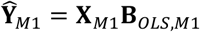 and 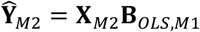, where *B*_*OLSM1*_ are the least-squares solution for **X**_*M*1_ and **Y**_*M*1_, and then calculated the correlation between 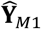 and 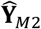 (Figure S4H).

### Statistical Analyses

Data analysis was performed using custom routines in Python, MATLAB (MathWorks) and GraphPad Prism 10 (GraphPad Software). No specific analysis was used to estimate minimal population sample or group size, but the number of animals, sessions and recorded cells were larger or similar to those employed in previous related work ^1,31,62,73,89,113^. Unless otherwise noted, non-parametric Wilcoxon rank-sum or Wilcoxon signed-rank test was used for unpaired and paired data comparisons respectively, and ANOVA and Kruskal-Wallis was used for multiple comparisons. All statistical tests were two-tailed with *p* < 0.05 as the cutoff for statistical significance, which is indicated by asterisks (\**p*<0.05, \*\**p*<0.01, \*\*\**p*<0.001, and \*\*\*\**p*<0.0001). Error bars show SEMs, and boxplots show median (central mark), 75th (box), and 90th (whiskers) percentile, unless indicated otherwise. For boxplots, points below the 10th percentile and above the 90th percentile are defined as outliers; outliers are not displayed in the plots but included in statistical analysis. Due to experimental constraints of optogenetic experiments, experimenters were not blind to these manipulations.

