## Supplemental figures for "A hippocampal population code for rapid generalization"

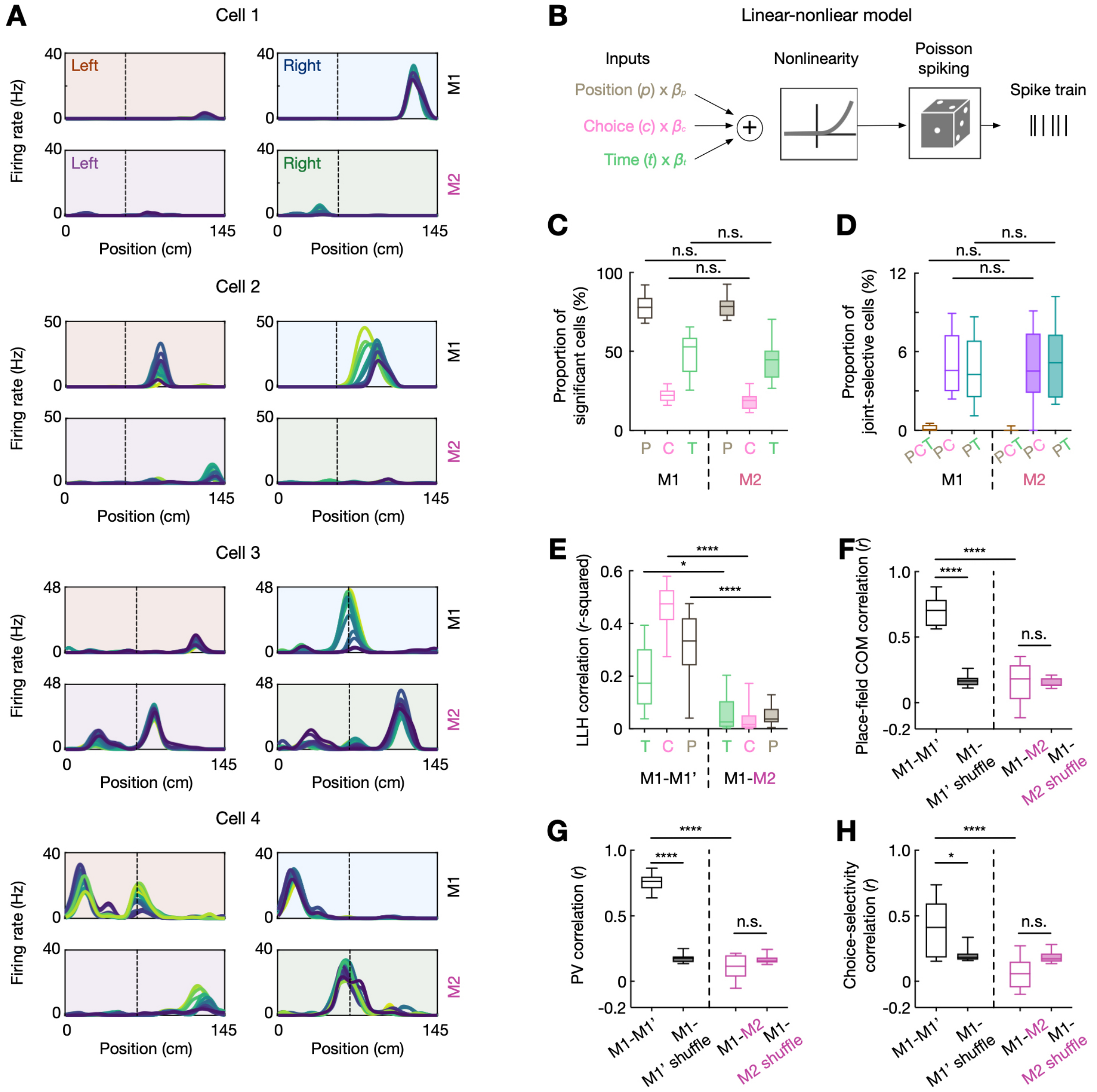


**Figure S1. Quantifying CA1 single-cell coding properties.** Related to Figure 1.
**(A)** Examples of single-cell coding of position, time and choice. Data are presented as in Figure 1D. Cell 1 shows a typical place coding cell, with a single stable place field on the R-trajectory of M1. Cell 2 shows drifting place fields on M1, with a typical backward shift of the R-side place field over trials. Cell 3 shows a drifting place field on R-trajectory of M1, but stable place fields on M2. Cell 4 shows conjunctive coding.
**(B)** Schematic of linear-nonlinear (LN) model framework for quantifying single-cell coding properties [96]. Position (P), time (T) and choice (C) were linearly combined with a vector of learned parameters ($\beta$), and then multiplied by an exponential nonlinear function, which accounts for the response nonlinearities such as rectification and saturation, followed by Poisson spike generation.
**(C)** Proportion of cells with significant selectivity to one of the three variables (P, T, and C) on M1 and M2. Significance was assessed via 10-fold cross-validation. As expected, the majority of CA1 cells exhibited selectivity for spatial locations (P). In addition, a

substantial number of CA1 cells showed selectivity for choice (C) and time (T). The proportions of selective cells were similar across two mazes (n.s., $p$ = 0.99, 0.96, and 0.96 for P, C, and T, respectively, one-way ANOVA with Tukey's *post hoc*).
**(D)** Proportion of cells that showed joint selectivity for more than one variable (i.e., mixed selectivity; similar proportions across two mazes, n.s., all $p > 0.99$, one-way ANOVA with Tukey's *post hoc*).
**(E)** Correlation of single-cell selectivity across mazes. Relative contribution of time, choice, position in predicting the single-cell spikes, quantified by the log-likelihood (LLH), was computed and compared across two sessions (M1 vs. M1', and M1 vs. M2). While the contributions were largely consistent across two sessions on the same maze, such selectivity did not generalize across two different mazes (****$p$(P) = 1.40e-7, *$p$(T) = 0.01, ****$p$(C) = 1.13e-11, one-way ANOVA with Tukey's *post hoc*).
**(F)** Single-cell place-field center-of-mass (COM) shift across M1 and M1' and across M1 and M2 (****$p$ < 8.64e-13, ****$p$ < 1e-16, n.s. $p$ > 0.99, one-way ANOVA with Tukey's *post hoc*).
**(G)** Population vector (PV) correlation across M1 and M1' and across M1 and M2 (****$p$ < 1e-16, n.s. $p$ > 0.99, one-way ANOVA with Tukey's *post hoc*).
**(H)** Choice selectivity correlation across M1 and M1' and across M1 and M2 (*$p$ < 0.012, ****$p$ < 4.44e-6, n.s. $p$ = 0.11, one-way ANOVA with Tukey's *post hoc*).
Shuffles in **(F-H)**: cell-identity shuffles.

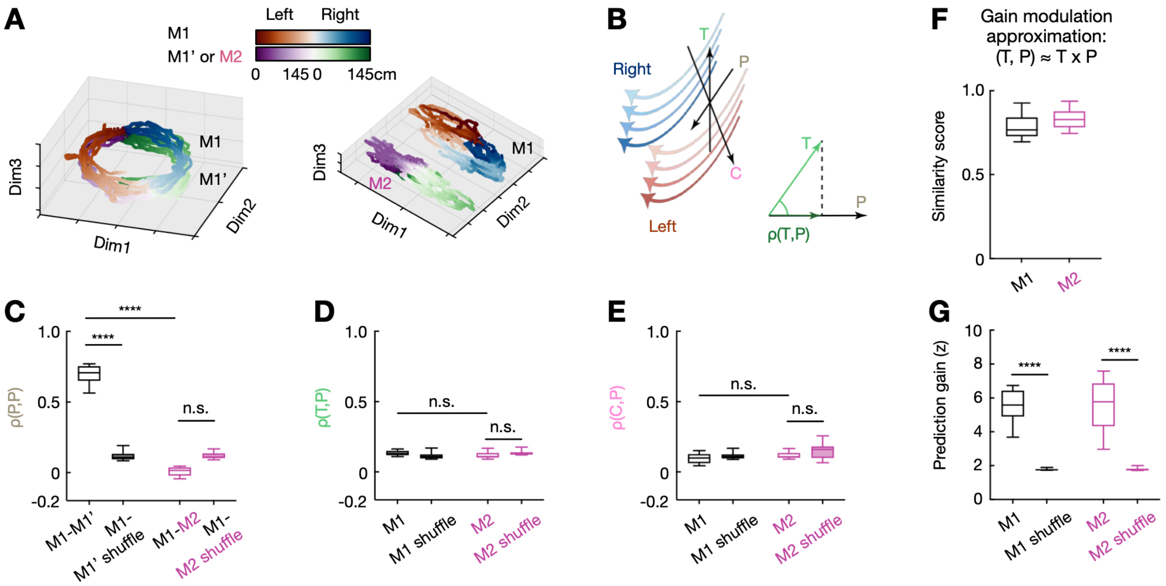


**Figure S2. Additional analysis quantifying orthogonal coding subspaces for position, time and choice.** Related to Figure 1.

**(A)** Different views of the neural manifolds shown in Figure 1F (*left*) and Figure 1G (*right*).

**(B)** Schematic for the method used to quantify the subspace overlap using the direction of changes in neural dynamics [101]. Across trials (T) and choice types (C), the neural activity associated with a particular physical location (P) changed (*left*). We assessed the alignment (*ρ*; *right*) between the directions of these changes (arrowheaded lines for T and C) and the position coding direction (arrowheaded line for P), such that *ρ* = 0 is for randomly oriented vectors (i.e., orthogonal), and *ρ* = 1 is for a perfect alignment (i.e., parallel).

**(C-E),** Alignment (*ρ*) of the position coding directions across two sessions **(C)**, and *ρ*(T, P) **(D)** and *ρ*(C, P) **(E)** within the same session (*****p* = 4.71e-13, n.s., *p*'s > 0.05, one-way ANOVA with Tukey's *post hoc*). Shuffles: cell-identity shuffles.

**(F and G)** An alternative method to assess subspace overlap using gain modulation approximation [55]. We assessed how well the joint population dynamics (T, P) can be approximated by multiplicative dynamics of T and P (T x P; i.e., factorization).

**(F),** High similarity score (ranging from 0 to 1) [55] between the joint model (T, P) and the gain modulation approximation (T x P), supporting a factorized (disentangled) representation of time and space.

**(G)** High prediction gain using the gain modulation approximation (z-scored based on mean and SD of the shuffled distribution; ****p* = 1.58e-10, *n* = 32 and 12 session pairs for M1-M1' and M1-M2, respectively, and one-way ANOVA with Tukey's *post hoc*). Shuffles: trial-time label shuffles.

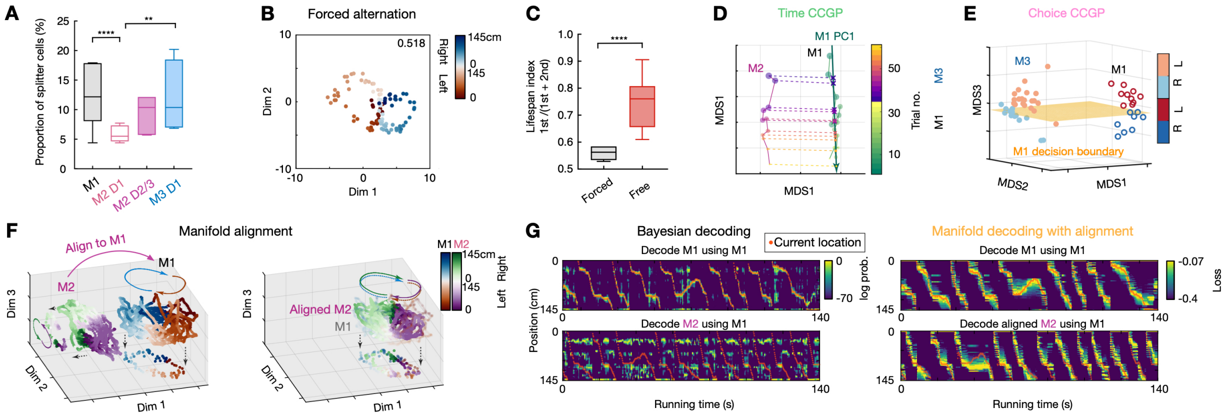

**Figure S3. Additional examples and analysis for generalizable coding dimensions in the hippocampus.** Related to Figure 2.

**(A)** Proportion of splitter cells across different mazes (****$p$ = 3.57e-5, **$p$ = 0.00072, z-test for proportions).

**(B)** Visualization of the topological structure of the neural manifold during the forced alternation version of the task on M1. Note the 2 loops (lifespan index = 0.518).

**(C)** Lifespan index of manifold topology during forced versus free (memory-guided) alternation tasks (****$p$ = 1.17e5, $n$ = 7 forced and 15 free alternation sessions from 5 mice, rank-sum test).

**(D)** Illustration of CCGP for time in one representative session. The first principal component (PC1) of the M1 data was used as a decoder to decode the trial time of the M2 data. Dashed lines: trials from M2 projected onto M1 PC1. Note the increasing projected values over trial time, yielding a good CCGP.

**(E)** Illustration of CCGP for choice in the same example session. A linear decoder (yellow plane) was trained to differentiate between L and R choice on M1. If the choice is represented in an abstract and generalizable format, the decoder will be able to decode choices on a different maze (M3), yielding a good CCGP for choice, as it was the case.

**(F)** Example of M1 and M2 manifold alignment using Procrustes transformation.

**(G)** Decoding example using classical Bayesian decoding (*left*) and manifold decoding after00 alignment (*right*). The decoders were built on M1 data to decode position of M2 data. Red dot: animal's current location. Only running periods were used.

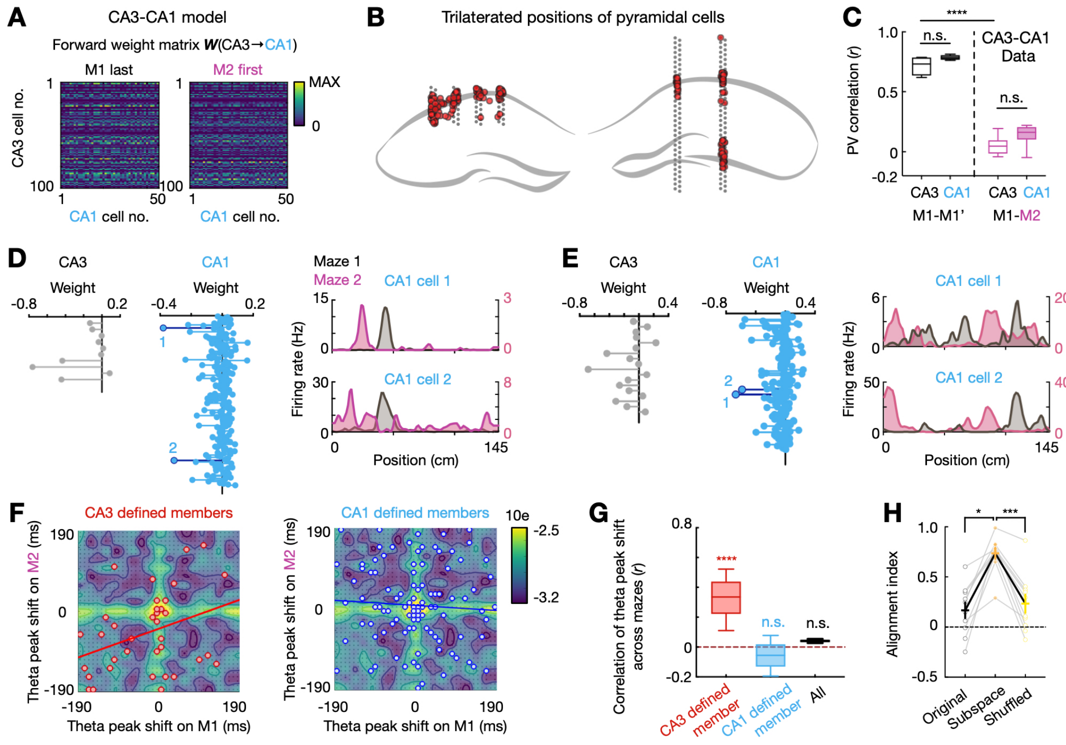

**Figure S4. Additional examples and analysis for coherent hippocampal remapping.** Related to Figure 3.

**(A)** Example feedforward CA3-CA1 weight matrix (**W**) during the last trial on M1 and first trial on M2. Although the spatial input abruptly changed across the two trials, the structure of the weight matrix was relatively stable (subtle changes from synaptic noise over time).

**(B)** Schematic of an example dual site recording in CA1 and CA3. The trilaterated positions of recorded cell somas are shown. Contours of the pyramidal layers are traced from the mouse brain atlas [114].

**(C)** In experimental data, both CA3 and CA1 showed global remapping across M1 and M2. Population vector (PV) correlation is shown (****$p$ = 1.04e-13 and 1.10e-13 for CA3 and CA1, respectively, $p$ = 0.054 and 0.64 for M1-M1' and M1-M2, respectively, one-way ANOVA with Tukey's *post hoc*).

**(D** and **E)** Examples of CA3-CA1 assemblies from experimental data. Data are shown as in Figures 3G and 3H.

**(F)** Temporal shift of theta covariance peak of CA1 cell pairs on M1 versus M2. Note that the distance between two place fields are correlated with the temporal shift of spikes within theta cycles between the two place cells [104,111,112], and thus the temporal shift served as an alternative remapping measure to Figures 3I and 3J. Assembly member pairs detected using the CA3-CA1 coactivation matrix are shown on the top (red dots), and those using the CA1-CA1 coactivation matrix are shown on the bottom (blue dots). Data are presented as in Figure 3I.

**(G)** Correlation of theta peak shift across mazes for CA3 defined CA1 member pairs, CA1 defined CA1 member pairs, and all CA1 pairs (**** $p$ < 1e-16, n.s., $p$ = 0.31 for CA3 versus CA1 defined member pairs, rank-sum test compared to 0).

**(H)** Communication subspace alignment with Reduced Rank Regression (see Methods). The alignment was significantly higher than that from their temporally shuffled data (***$p$ = 0.0004), and from the alignment without being reduced to the communication subspace (*$p$ = 0.011, $n$ = 10 session pairs from 3 mice, Friedman test with Dunn's *post hoc*).

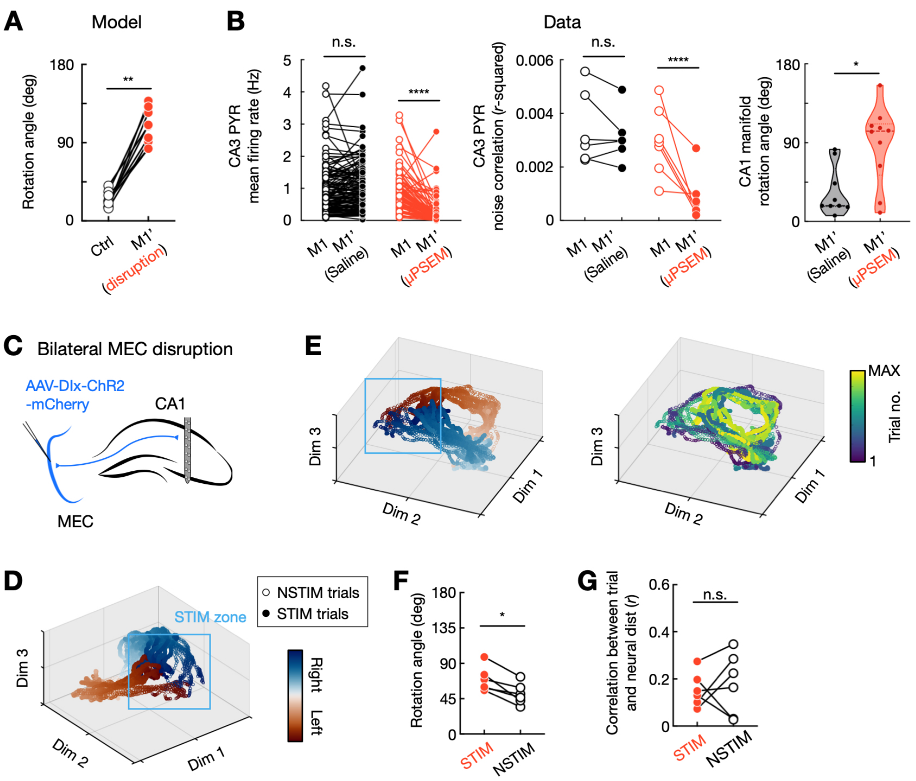


**Figure S5. Additional analysis for CA3 disruption and bilateral MEC disruption.** Related to Figure 4.

**(A)** Rotation angle between control (M1) and input disruption (M1') trial blocks from 9 model simulations (***p* = 0.0039, Wilcoxon paired test).

**(B)** *Left and middle*, Reduced firing rate (*left*; n.s., *p* = 0.21, *****p* = 4.86e-18, *n* = 128 and 83 cells, rank-sum test) and reduced noise correlation (*middle*; n.s., *p* = 0.44, **p* = 0.016, *n* = 6 and 7 session pairs, Wilcoxon paired test) of experimentally recorded CA3 pyramidal cells during chemogenetic manipulation, consistent with the model manipulation (Figure 4A). *Right,* Rotation angle of CA1 neural manifolds between the first and second session calculated for the control and CA3 disruption conditions (**p* = 0.01, *n* = 10 control and 9 CA3 disruption session pairs from 4 mice, rank-sum test).

**(C)** Schematic of the approach used for bilateral MEC disruption. AAV-Dlx-ChR2 was injected bilaterally into MEC to express ChR2 in all GABAergic cells [73,103], and optical fibers were implanted bilaterally over MEC. Optogenetic stimulation was delivered at a segment of the maze in a block of 10 trials (STIM) interleaved with 10-trial blocks of no stimulation (NSTIM).

**(D)** Manifold dynamics during STIM (solid circles) and NSTIM (hollow circles) trials, color coded by positions.

**(E)** A different view of the neural manifold shown in **(D)**, color coded by positions (*left*) and trial time (*right*). Note the off-manifold remapping at the STIM zone.

**(F),** Rotation angles between STIM-NSTIM trials and NSTIM-NSTIM trials (**p* = 0.031, *n* = 6 sessions from 3 mice, Wilcoxon paired test).

**(G)** MEC perturbation did not affect CA1 temporal representational drift with time (n.s., *p* = 0.69, *n* = 6 sessions from 3 mice, Wilcoxon paired test). Correlation between trial time and neural distances is shown.

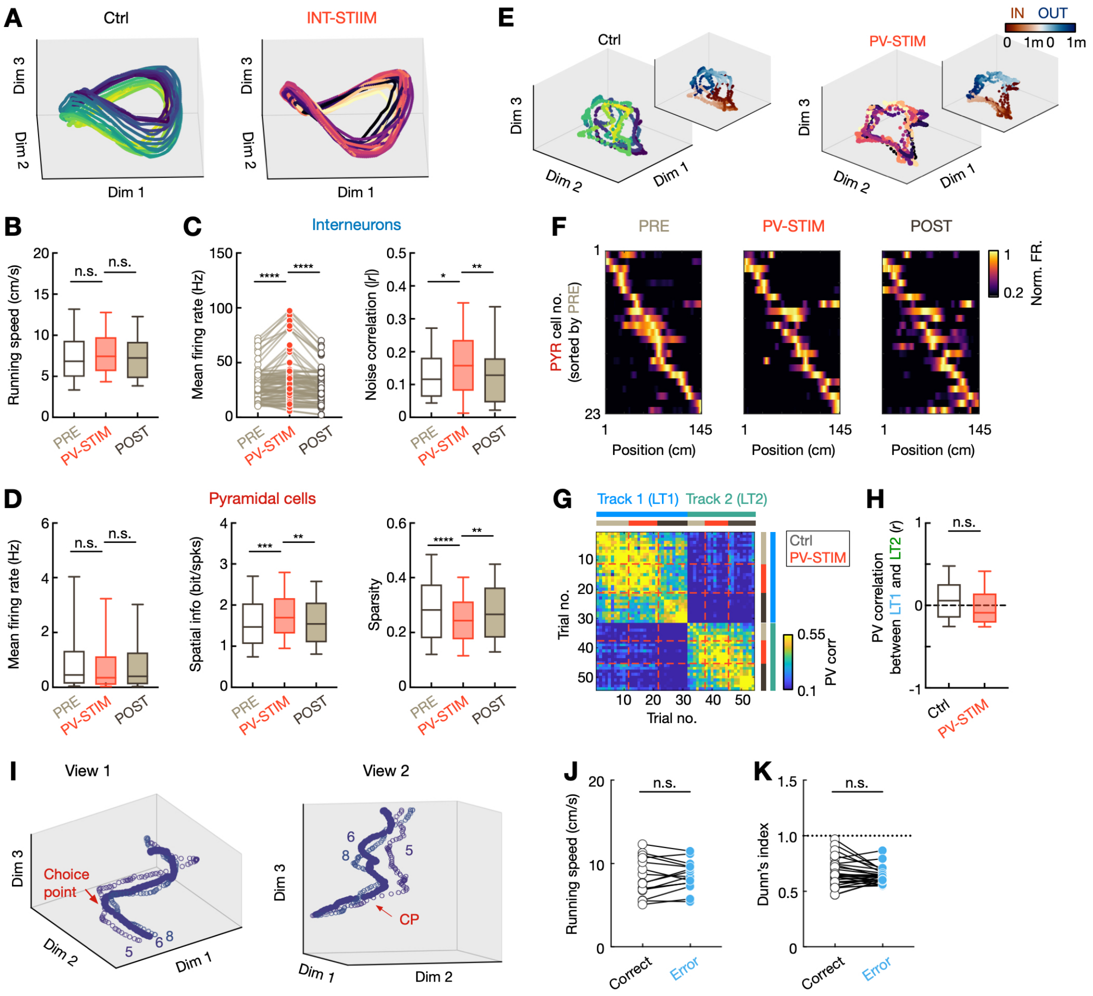


**Figure S6: Additional analysis for local interneuron manipulation.** Related to Figure 5.

**(A)** Neural manifolds from an example CA1 model simulation for the Ctrl and INT-STIM conditions, color coded by trial number. Note the ordinal drift in Ctrl but not INT-STIM.

**(B)** Running speed for PRE, PV-STIM, and POST trial blocks (n.s., $p$ = 0.13 and 0.058, $n$ = 9 sessions from 4 mice, Wilcoxon paired test).

**(C)** Increased firing rate (*left*; ****$p$ = 3.53e-6 and 1.74e-9, $n$ = 85 cells from 4 mice) and enhanced noise correlation (*right*; *$p$ = 0.0123, **$p$ = 0.0096, $n$ = 38 trial pairs from 4 mice, repeated-measures one-way ANOVA with Tukey's *post hoc*) of CA1 interneurons with the manipulation.

**(D-F)** Preserved place-field properties during optogenetic PV manipulation ($n$ = 9 sessions from 4 mice).

**(D)** *Left*, mean firing rate of CA1 PYRs during PRE, PV-STIM, and POST trials (n.s., $p$ = 0.53 and 0.94, one-way ANOVA with Tukey's *post hoc*). *Middle*, Enhanced spatial information of CA1 PYRs during PV-STIM trials (***$p$ = 0.0001, **$p$ = 0.0013, one-way ANOVA with Tukey's *post hoc*). *Middle*, Sparser place response of CA1 PYRs during PV-STIM trials (***$p$ = 4.67e-6, **$p$ = 0.002, one-way ANOVA with Tukey's *post hoc*; sparsity = 1 implies no sparseness).

**(E)** Neural manifolds for the Ctrl and PV-STIM condition from experimental data, color coded by trial numbers (insertions on the left show the same manifolds but color coded by locations and choice types).
**(F)** Normalized rate maps of all CA1 spatial-tuned pyramidal neurons during PRE, PV-STIM, and POST trials, sorted by peak activity during PRE trials, from an example session. Note consistent place fields across conditions.
**(G** and **H)** Optogenetic PV manipulation did not affect remapping across different linear tracks (LT1 and LT2). Population vector correlations between LT1 and LT2 Ctrl (PRE and POST; light gray, PRE; dark gray, POST) and PV-STIM trials are shown (n.s., $p = 0.11$, rank-sum test compared to 0).
**(I)** Two additional views of the error trial shown Figure 5J.
**(J** and **K)** Control analysis for error trials of the delayed alternation task.
**(J)** Running speed for correct and error trials (n.s., $p = 0.42$, $n = 15$ sessions from 5 mice, Wilcoxon paired test; sessions with less than 3 error trials were excluded).
**(K)** Error trials were on-manifold. Dunn's index evaluating the distance of neural trajectories of correct and error trials to the neural manifold (n.s., $p = 0.36$, $n = 28$ sessions from 5 mice, Wilcoxon paired test)
